# BoviScan: a Phage ImmunoPrecipitation Sequencing library to characterize the antibody response against *Mycoplasma bovis*

**DOI:** 10.64898/2026.09.14.751406

**Authors:** Yonathan Arfi, Eric Baranowski, Carole Lartigue, Gabriela Pretre, Christine Citti, Pascal Sirand-Pugnet

## Abstract

*Mycoplasma bovis* is a major cattle pathogen and a global concern due to the limited efficient prophylactic tools and the emergence of antibiotic-resistant strains. Current control strategies rely primarily on surveillance, diagnostic and segregation or culling of infected animals. Despite substantial efforts to develop vaccines against *M. bovis*, no broadly effective solution is currently available. This situation is partly attributable to important gaps in our understanding of the host humoral immune response following infection, vaccination, or challenge. ELISA and Western blot assays have been widely used to assess *M. bovis*-specific antibody responses. While robust and easy to implement, these approaches are difficult to scale for hundreds of antigens in large animal cohorts and provide limited information on the specific antigens and epitopes targeted by the antibody response. To address these limitations, we developed BoviScan, a Phage Immunoprecipitation Sequencing (PhIP-Seq) library covering the pan-proteome of 295 *M. bovis* strains. We describe the design, synthesis, and validation of BoviScan using a panel of known-positive sera. We then applied the library to characterize antibody responses in a small cohort of experimentally immunized cattle and to monitor the longitudinal evolution of the humoral response in a single animal. Our results show that BoviScan enables high-throughput, high-resolution mapping of the antibody responses against *M. bovis*. This approach could be highly valuable for the study of host-pathogen interactions in mycoplasmas and the development of improved vaccines and serological assays.

## Introduction

*Mycoplasma bovis* is a bacterium belonging to the class *Mollicutes*, and is characterized by the lack of cell wall and a reduced genome (∼0.9 Mb)^1,2^. It is able to colonize the mucosa of a number of domesticated and wild ruminant species, including bison and most cattle^3–5^. Colonization of the host can be asymptomatic but often transition to pathogenicity. *M. bovis* is associated with, and considered to cause, a range of chronic, low mortality and high morbidity diseases, including pneumonia, bronchitis, mastitis and arthritis^6,7^. In addition, it is also found frequently associated with other pathogens, and is considered a major contributor to the Bovine Respiratory Disease Complex (BRDC)^8^. Transmission occurs through contact between animals, but also with fomites, since *M. bovis* can survive for extended periods of time on inert surfaces in the farms^9^.

While the pathogenicity of *M. bovis* is not yet fully understood, a number of studies have highlighted a particularly wide array of persistence mechanisms and effectors^10^. These include surface antigen phase variation through ON-OFF switching in expression of variable, surface lipoproteins^11,12^; the production of biofilm^13^; enzymatic degradation of neutrophil extracellular traps DNA^14,15^; and direct targeting and degradation of host antibodies through the MIB-MIP system^16,17^. These mechanisms enable *M. bovis* to evade both the cellular and humoral component of the host immune response, and to establish chronic infections^18^.

Due to its strong negative impact on animal health, welfare and productivity, *M. bovis* is recognised as a major issue for both the dairy and meat cattle industry. Prevention currently relies mainly on diagnostics, sanitary control measures, and antimicrobial therapy. However, *M. bovis* is naturally resistant to molecules targeting the cell-wall, and the recent emergence of resistance to the few available antibiotics further limits the treatment options^19,20^.

Initially identified in the USA in the early 1960’s, *M. bovis* is now found worldwide^21^. A severe outbreak occurred in 2017 in New Zealand, a country initially free of this pathogen^22,23^, and resulted in the culling of 126,000 cattle. The outbreak is suspected to have originated from contaminated, imported bovine semen used for artificial insemination, and following this single introduction event to have rapidly spread throughout the country. Owing to the lack of efficient prevention strategy at the time, an ambitious national eradication programme based on pathogen detection and herd culling was implemented over the years. This eradication effort came at extremely high economic and social cost^24,25^.

While substantial efforts have been made toward the development of effective vaccines^26,27^, major gaps remain in our understanding, in particular of the host humoral immune response during *M. bovis* infection or vaccination. Indeed, currently two main tools are used to characterize the antibody response: ELISA, and Western Blot. Both approaches can be performed to probe various antigens, ranging from simple peptides to recombinant proteins, and up to whole bacterial cell extracts. In the case of cell extracts, the whole proteome of a strain can be probed, but information on which proteins and which epitopes are targeted is impossible to obtain with ELISA, while Western Blot can be more informative when coupled to 2D gel electrophoresis^28^. Conversely, using a specific protein as target can be highly informative, but costly and hard to scale up enough to interrogate the whole proteome or the pan-proteome of multiple strains.

Recently, a new high-throughput strategy has been developed to characterize the composition of the antibody pool targeting the proteome of a set of pathogens. This method is called Phage Immuno-Precipitation Sequencing (PhIP-Seq)^29–31^. It relies on the production of an oligonucleotide library that encodes peptides covering the whole proteome of the target species. This peptide library is cloned as a bacteriophage display library, which is then immuno-precipitated using the antibodies contained in a sample. The peptide-encoding DNA contained in the immune-precipitated phages is subsequently amplified by PCR and analysed by high-throughput DNA sequencing. The number of reads mapped for each peptide coding sequence is correlated to relative enrichment of the corresponding peptide, and to the presence of the cognate antibody in the sample (Figure 1). Therefore, PhIP-Seq allows the characterization of the antibody repertoire in a high-throughput and semi-quantitative manner, and to the single epitope level.

**Figure 1:**
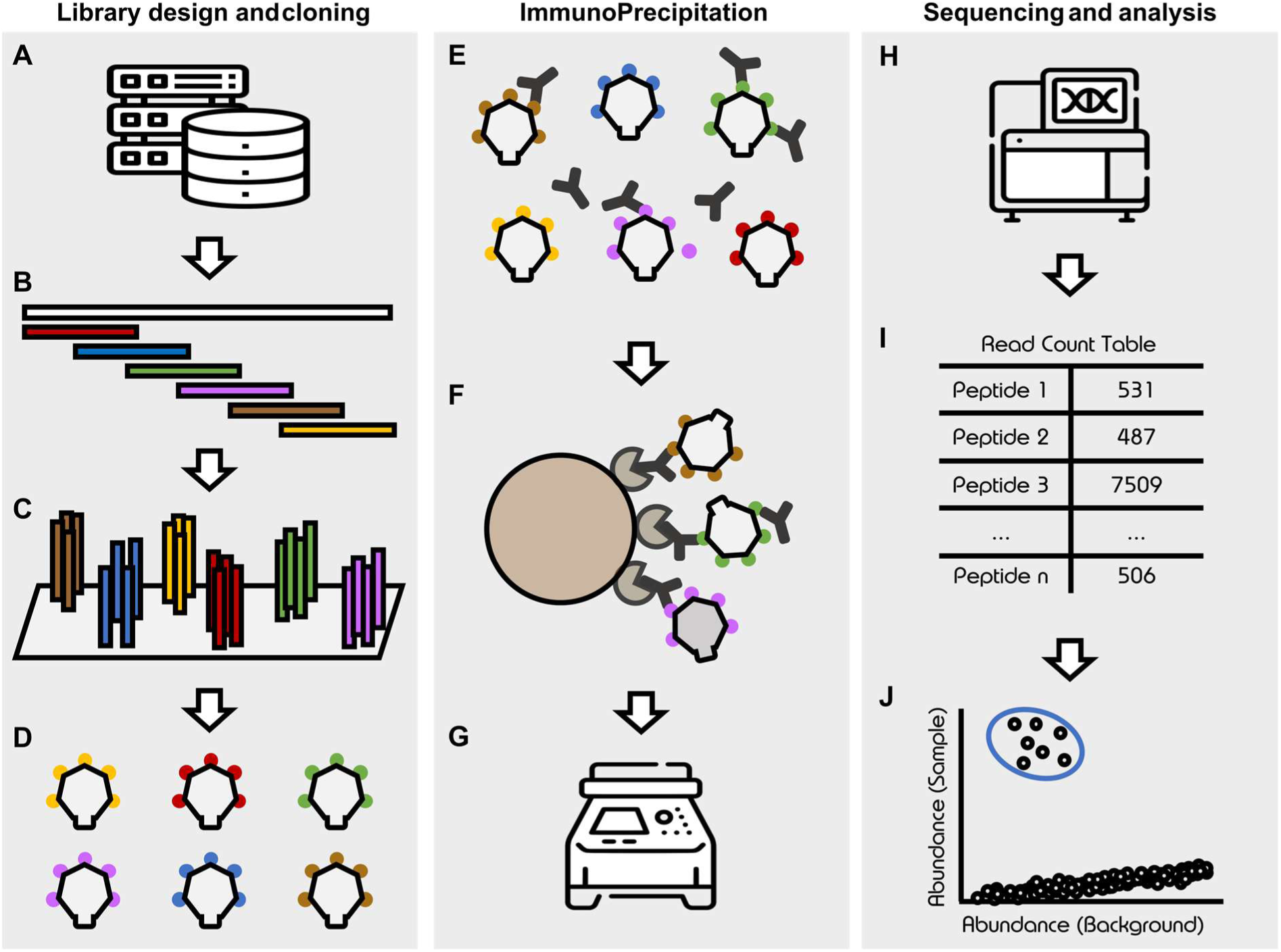
Overview of the PhIP-Seq method. A. Protein sequences are collected from various databases. B. Each protein sequence is split into a set of overlapping peptide sequences. C. An oligonucleotide pool encoding all the peptides sequences is synthesized. D. The amplified oligo-pool is cloned as a bacteriophage display library. E. The bacteriophage library is incubated with an antibody-containing sample. F. Magnetic beads coated with immunoglobulin-binding proteins are used to pull down the antibodies and their bound phages. G. PCR is performed on the DNA of precipitated phages to amplify the peptide-coding region. H. The amplicons are sequenced by high throughput sequencing. I. The reads are mapped to the library’s reference sequences to obtain a count table. J. Statistical analysis of the count table is performed to determine peptide enrichment compare to a background control.

This method was initially developed using a library covering a part of the human proteome, in order to probe the human antibody repertoire against human epitopes and assist in the discovery of new autoantigens^32^. It was then applied with great success to perform a comprehensive serological profiling of human sample against a wide array of viruses, using the VirScan library which encoded the pan-proteome of 206 species of virus (and over 1000 different strains)^33^. Over the years, multiple PhIP-Seq libraries have been developed, targeting a diverse range of epitopes, including the mouse proteome^34^, the human gut microbiome^35^, known-allergens from food and environmental origin^36^, known toxins or virulence factors^37^. Some libraries were also designed to target specific pathogens, including SARS-Cov2^38^ *Plasmodium falciparum*^39^, *Trypanosoma cruzi*^40^, *Trypanosoma brucei*^41^, and *Schistosoma mansoni*^42^. However, there is currently no published library focusing on a given bacterial pathogen.

In order to evaluate the potential benefits of this method to study the humoral response to *M. bovis* in bovines, we have developed “BoviScan”. This PhIP-Seq library is comprised of 37,285 peptides representing 421,793 CDSs gathered from the genomes of 295 *M. bovis* strains. After cloning and quality controls, the BoviScan library was assessed using known positive samples; then used to compare the antibody response across a small animal cohort, and to perform a longitudinal study of the response in a single animal.

## Material and methods

### PhIP-Seq library design

Protein sequences used for the design of BoviScan were obtained from the NCBI Genomes database (https://www.ncbi.nlm.nih.gov/datasets/genome/) on 24/01/2023. The protein-coding sequences from 557 annotated assemblies were downloaded. When available, both the NCBI RefSeq annotation and the GenBank submitter annotation were collected. An additional 9 entries were manually added to the dataset: one corresponding to *M. bovis* strain KG4397 isolated in Japan^43^, one to the atypical strain JTCL1 isolated from rabbit^44^ and six corresponding to strains isolated in Israel and deposited in the NCBI Genomes database but unannotated. For these six strains, annotation was performed using *prokka*^45^ (version 1.14.6 on a Galaxy webserver (https://usegalaxy.eu)^46^.

All the CDSs were collected and used to generate a library of encoded peptides. Each CDS was given a unique identifier comprised of its Assembly Accession number coupled to Protein identifier. This step was necessary because the RefSeq annotated CDSs can sometimes share a Protein identifier.

In order to split each protein into overlapping peptides (or “tiles”), we then used the *pepsyn* package (https://github.com/lasersonlab/pepsyn), following the blueprint provided by Mohan *et al.* 2018 ^29^. All the CDSs were first pooled in a single file and then broken down in 56 amino-acids long sequences overlapping each other by 28 amino-acids (arguments: *pepsyn x2ggsg - - | pepsyn tile -l 56 -p 28 - - | pepsyn disambiguateaa - -*). In addition, for each CDS a 56 amino-acids long tile covering the C-terminal portion of the sequence was generated (arguments: *pepsyn x2ggsg - - | pepsyn ctermpep -l 56 --add-stop - - | pepsyn disambiguateaa - -*).

The peptides contained in the two corresponding output files (ORF tiles and the Cter tiles) were then clustered using CD-HIT^47^ at 98% sequence identity (arguments: *cd-hit -i orf_tiles -o clustered98 _orf_tiles -c 0.98 -G 0 -A 50 -M 0 -T 1 -d 0*; and *cd-hit -i cter_tiles -o clustered98_ cter_tiles -c 0.98 -G 0 -aL 1.0 -aS 1.0 -M 0 -T 1 -d 0)*. The clustered ORF tiles and Cter tiles were then pooled in a single file, and when needed their size padded to 56 amino-acids long using pepsyn (argument: *pepsyn pad -l 56 --c-term - -*).

The tiles amino-acid sequences were then reverse translated into DNA sequences with pepsyn, using the *Escherichia coli* codon usage frequency table to guide the choice of codons, removing any EcoRI and HindIII restriction site. In addition, each tile coding sequence was appended with a common prefix (5’-AGGAATTCCGATGCG-3’) and suffix (5’-GCCTGGAGAAAGCTTCG-3’) in order to enable its cloning downstream (argument: *pepsyn revtrans --codon-freq-threshold 0.01 --amber-only - - | pepsyn prefix -p AGGAATTCCGATGCG - - | pepsyn suffix -s GCCTGGAGAAAGCTTCG - - | pepsyn recodesite --site EcoRI --site HindIII --clip-left 15 --clip-right 17 --codon-freq-threshold 0.01 --amber- only - -*).

The overall output of this process is a fasta file containing the coding sequences of all the peptide tiles; with each sequence being 200 nucleotides long (15 nucleotides prefix, 168 nucleotides peptide CDS and 17 nucleotides suffix).

The same process was used to design the control tiles. We first collected the coding sequences of the four proteins forming the viral particle of the Foot-and-Mouth Disease Virus (FMDV) type O (PDB: 7ENP). As FMD is a highly controlled disease, seronegative samples should be prevalent while seropositive control samples should be readily available. Overall, 43 FMDV tiles were generated (39 ORF tiles and 4 C-Terminal tiles) using the same tiling logic as above. We then elected to add to the library a set of five well-defined epitope-tags, namely HA tag (YPYDVPDYA), c-Myc tag (EQKLISEEDL), Avi-tag (GLNDIFEAQKIEWHE), StrepTag ii (WSHPQFEK) and ALFA tag (SRLEEELRRRLTE). Each tag sequence was added to the C-Terminal end of each FMDV tile via a (GS)_3_ linker. In order to maintain the 56 amino-acids length, part of the FMDV portion of the tagged-tiles was trimmed from the N-terminal side. Overall, 215 FMDV-Tag tiles were generated and appended to the BoviScan design.

All the sequences described above are available as supplementary material.

### PhIP-Seq library cloning

The designed peptide library was synthesized as an oligo-pool (Twist Bioscience). The single stranded DNA mix was resuspended in TE buffer (Tris 10 mM, EDTA 0.1 mM, pH 8) to a final concentration of 10 ng.µL^-1^. Double stranded DNA was generated by PCR, using the ssDNA oligo pool as template. To do so, PCR was performed using the Kapa HiFi Hotstart kit (Roche) and primers matching the prefix and suffix (BoviScan_Prefix-F: 5’-AGGAATTCCGATGCG-3’ and BoviScan_Suffix-R: 5’-CGAAGCTTTCTCCAGGC-3’). PCR was performed following the manufacturer’s recommendations: all reactions were performed in a total volume of 25 µL, Kapa Fidelity Buffer was used at 1X, Kapa dNTPs were used at 0.3 mM, primers were used at 0.3 µM, Kapa Hifi polymerase was used at 0.5 Unit per reaction, and 1 ng of ssDNA template was used. The thermal cycling conditions were as follows: 95°C 3 min; 16 cycles of 98°C 20 s, 62° 20 s, 72°C 15 s; 72° 1 min; 10°C hold. Low cycle numbers were used to limit the occurrence of potential replication error. A set of 32 PCR reactions were performed and pooled.

The PCR quality was checked by agarose gel electrophoresis. We noted that the amplicons generated had a higher apparent size than expected. After investigation, we identified that this higher apparent size was an electrophoresis artefact caused by so-called “PCR bubble products”. These “bubble products” are heteroduplexes formed during PCR, due to the partially heterogeneous nature of the amplified library, are well described in the Illumina literature and are not problematic for downstream applications. ONT sequencing of these amplicons confirmed that their true size was correct, and the amplicons were subsequently purified using AMPure XP magnetic beads (Beckman) and eluted in water. DNA concentration was checked using a Qubit fluorometer and the associated Qubit dsDNA Broad Range reagents (Thermo Scientific).

The purified amplicons were digested using the restriction enzymes *Eco*RI-HF and *Hind*III-HF (NEB) following the manufacturer’s protocol. Five independent reactions were set up, each in a final volume of 50 µL, containing CutSmart buffer at 1X final concentration, 1 µg of DNA and 20 Units of both enzymes. Restriction reactions were carried out at 37°C for 1 hour. Complete digestion of the DNA was assessed by agarose gel electrophoresis, and the digested DNA was subsequently pooled and purified using AMPure XP magnetic beads (Beckman) and eluted in water. DNA concentration was checked using a Qubit fluorometer and the associated Qubit dsDNA Broad Range reagents (Thermo Scientific).

Cloning of the DNA library in T7 bacteriophages was performed using the T7Select 10-3 cloning kit (Merck) following the kit’s instructions. The digested DNA was first ligated into the provided T7Select *Eco*RI/*Hind*III vector Arms. To do so, ligation was performed in a final volume of 5 µL, using 500 ng of T7 vector Arms, 8 ng of insert DNA, 5 Units of T4 DNA ligase (Thermo Scientific) and 0.5 µL of 10X ligase buffer (Thermo Scientific). Ligation was carried out at 16°C in a thermocycler for 16 hours.

The reconstituted T7 genomes were then rescued by *in vitro* packaging. To do so, 1 µL of ligation reaction was added to 5 µL of T7Select Packaging Extract (Merck) and incubated at 23°C for 2 hours. Packaging was stopped by addition of 56 µL of LB media and the mix was stored on ice.

The packaging reaction was checked by performing a phage titration assay. Briefly: *E. coli* BLT5403 were grown in liquid M9TB medium, supplemented with 100 µg.mL^-1^ of ampicillin until the OD_600 nm_ reached ∼1. A sample of the packaging reaction was serially diluted in LB in 1/10^th^ increment down to a 10^-7^ dilution. The number of phages in 100 µL each dilution was obtained by mixing with 250 µL of the *E. coli* BLT5403 and 3 mL of warm Top agar and plating on LB agar plates supplemented with 100 µg.mL^-1^ of ampicillin. After 4 hours of incubation at 37°C, the number of plaques forming units (PFUs) were counted on each plate, to estimate the phages titre in the packaging reaction.

The rescued library was then amplified using the recommended plate amplification method. To do so, *E. coli* BLT5403 were grown in liquid M9TB medium, supplemented with 100 µg.mL^-1^ of ampicillin, until the OD_600 nm_ reached ∼1. Samples of the stopped packaging reaction were mixed with *E. coli* cells in order to create a 0.001 multiplicity of infection (1x10^6^ phages per 1x10^9^ bacteria). 1 mL of these phages-infected cells (corresponding to 1x10^8^ bacteria) were mixed with 10 mL of warm Top agar and plated on LB agar plates (in 140 mm round Petri dishes) supplemented with 100 µg.mL^-1^ of ampicillin. Plates were incubated at 37°C for 3-4 hours, until confluence was observed. In order to recover the phages, 10 mL of phage extraction buffer (Tris 20 mM, NaCl 100 mM, MgSO_4_ 6 mM, pH 8) were overlaid onto each plate, and incubated overnight at 4°C. The liquid was harvested the next days by tipping the plates and pipetting the buffer. The extracts from all the plates were combined, and centrifuged at 3000 rcf at 4°C for 5 min, in order to remove any large debris of agar or cells. The supernatant was carefully transferred in a new container and filtered twice – first on a 0.45 µm filter, then on a 0.22 µm filter. The resulting phage suspension was titrated by performing a phage titration assay (as described above) in duplicate. The phage suspension was then mixed with 0.1 volume of sterile glycerol, mixed, aliquoted and frozen at -80°C until use.

Quality control of the cloned library was performed by PCR on a sample of the library, coupled to high throughput sequencing of the amplicons. PCR was performed using the Kapa HiFi Hotstart kit (Roche), using the same reaction mixture described above. In this case, the tile-encoding locus in the T7 phages was amplified using primer T7SelectUp (5ʹ-GGAGCTGTCGTATTCCAGTC-3ʹ) and T7SelectDown (5ʹ-AACCCCTCAAGACCCGTTTA-3ʹ). Template consisted of 1 µL of T7 phage library, the number of PCR cycles was 25, and the annealing temperature was 66°C. All other parameters were unchanged. After agarose gel electrophoresis control, the amplicons were purified using AMPure XP magnetic beads (Beckman), eluted in water and concentration was checked using a Qubit fluorometer and the associated Qubit dsDNA Broad Range reagents (Thermo Scientific). Purified amplicons were submitted for sequencing (see details below).

### Phage immunoprecipitation

Phage immunoprecipitation was performed following a protocol derived from Mohan *et al*. 2018^29^ and Shrock *et al*. 2022^48^ First, for each sample to analyse, a 1.5 mL micro-centrifuge tube was blocked by incubation overnight at 4°C on a rotator with 800 µL of blocking buffer (Tris 20 mM, NaCl 137 mM, pH 7.5, Tween 20 0.1 % v/v, and BSA 3% m/v). The next day, the tubes were emptied by aspiration and kept at room temperature until use. Sera samples were prepared by diluting each serum at 1/100^th^ in Dulbecco’s Phosphate Buffered Saline (DPBS; Eurobio Scientific) and kept on ice. Monoclonal antibody samples were prepared by diluting 1 µg of IgG in 20 µL DPBS and kept on ice.

An aliquot of the frozen phage library was thawed on ice. For each sample, 3.7x10^9^ PFUs were diluted in a final volume of 1 mL of cold DPBS and stored on ice until use.

The antibody-antigen binding reaction was performed by mixing 20 µL of diluted serum (or monoclonal antibody) with 980 µL of phage library dilution, and incubating for 16 hours at 4°C on a rotator. For mock-IP controls, the 20 µL of sample were replaced by 20 µL of DPBS.

The next day, for each sample, 20 µL Protein G coupled magnetic beads (NEB) were prepared by removing the storage buffer, washing in DPBS, and resuspending in a final volume of 20 µL of DPBS.

Immunoprecipitation of the antibodies-phages complexes was carried out by adding 20 µL of washed Protein G magnetic beads to the 1 mL antibody-phage solution, and incubating at room temperature for 4 hours in a rotator. The magnetic beads were subsequently collected using a magnet, washed twice with 170 µL of TNP40 buffer (Tris 50 mM, NaCl 150 mM, pH 7.5, NP40 0.1% v/v) and once with 400 µL of DPBS. The beads were finally resuspended in 40 µL of water, transferred to a PCR tube, and heated to 70°C for 10 minutes. This bead suspension was stored at -20°C until use, or used immediately as template for PCR.

PCR was performed with the same protocol as for the quality control of the cloned library (see above), in a final volume of 50 µL and with 20 µL of the beads suspension as DNA template. After agarose gel electrophoresis control, the amplicons were purified using AMPure XP magnetic beads (Beckman), eluted in water and concentration was checked using a Qubit fluorometer and the associated Qubit dsDNA Broad Range reagents (Thermo Scientific). Purified amplicons were sent to sequencing (see details below).

### Sequencing

Sequencing library and sequencing itself were performed as a service by BGI Genomics. Library preparation was performed by using BGI Optimal DNA Library Prep Kit (BGI-Shenzhen, China). Purified amplicons were subjected the A-tailing reaction in order to add a single adenosine on the 3’ end of DNA. Subsequently, the library adaptors were connected to the two ends of DNA by ligation. The ligated products were amplified by PCR and subjected to quality control process. Next, the final double strand library products were denatured to generate the single stranded library product. Then, the circularization reaction was set up to generate single stranded circularized DNA products. Remaining single strand linear DNA was removed by digestion. The final single strand circularized library was amplified using the phi29 DNA polymerase and rolling circle amplification (RCA) to generate the DNA nano ball (DNB) which carries about 300 copies of the initial single stranded library molecule. The DNBs were then loaded into the patterned nanoarray and sequencing reads of PE150 bases length were generated using the DNBSEQ-G400 platform (BGI-Shenzhen, China).

### Sequencing data processing and analysis

All the sequencing data were processed using the Galaxy platform (https://usegalaxy.eu/). First, reads quality was assessed using Falco. Then, a read filtering, trimming and mapping pipeline was used to generate a read count table for each sample.

1. Reads trimming: *Trimmomatic* (version 0.39) was used in single-end mode and sequentially performs MINLEN 140 to drop any read shorter than 140 bases, AVGQUAL 28 to drop any read with average quality lower than 28, and HEADCROP 75 to remove the first 75 bases at the start of the read. This last step ensures the removal of sequences that are common to all reads and which correspond to T7 genome sequences and the prefix or suffix upstream of the tiles coding sequences.
2. The filtered and length-trimmed reads were then mapped to the reference library sequences using *Bowtie2* (version 2.5.5) in single-end mode, with an index built from the reference fasta file uploaded, and using default presets.
3. To generate a read count table, the .bam file outputted by the previous step was parsed using *Samtools coverage* (version 1.22.1), using the parameters: Minimum read length: 0; Minimum mapping quality: 0; Minimum base quality: 0; Required flags: none; Excluded flags: none.

The read count table provided the number of mapped reads for each tile present in the reference library, for each sample. This read count (RC) was then normalized to account for the differences in total reads across samples. To do so, each read count was divided by the total number of mapped reads (T) and multiplied by 10^6^, resulting in a Reads Per Million (RPM) value for each peptide (RPM = RC x 10^6^ / T).

Peptide enrichment analysis was performed by comparing the RPM values in samples to the values obtained in the mock-IPs. One method relied on EdgeR, ran through Galaxy using the *EdgeR-quasi* tool (quasi-likelihood pipeline) with parameters: Filter Low Counts: Yes; Minimum Count: 100; Filter on Total Count; Minimum log2 Fold Change: 2; P-Value Adjusted Threshold: 0.05; P-Value Adjustment Method: Benjamini-Hochberg; Use Robust Settings: Yes. The output of EdgeR is a list of statistically enriched peptides.

Alternatively, a Z-score was calculated for each peptide per sample, by subtracting the peptide’s mean RPM in the mock-IP from the peptide’s RPM value in the sample and dividing by the standard deviation of the peptide’s RPM in the mock-IP (Z-score = [RPM_sample_ – mean RPM_mock-IP_] / SD RPM_mock-IP_).

Statistical analyses and graphing were performed in Excel and Python (v3.12) using the libraries *pandas* (v3.0.2), *NumPy* (2.4.4) and *SciPy* (1.17.1), using Claude.ai (model Sonnet 4.6) to write and troubleshoot the corresponding scripts.

### Animal serum and monoclonal antibodies

All animal experiments were conducted in accordance with the European Community Council guidelines on the protection of animals used for scientific purposes (Directive 2010/63/EU) and were approved by the relevant national ethical authorities (APAFIS#4324-2016022418315195v2 and APAFIS#4672-2016030309521177v4; French Ministry of Agriculture, Ethics Committee no. 115).

Female BALB/c mice (8 weeks old; Janvier Labs) were housed in the animal facilities of the National Veterinary School of Toulouse (Ecole Nationale Vétérinaire de Toulouse, ENVT, Toulouse, France). Mice were immunized by subcutaneous injection (200 µL) with 20–40 µg of purified *M. bovis* recombinant proteins (Table 1) emulsified in complete Freund’s adjuvant on day 0, followed by booster injections formulated with incomplete Freund’s adjuvant on days 28 and 49. Sera were collected on day 71 by terminal cardiac puncture.

**Table 1.** *M. bovis* membrane-associated proteins produced in *E. coli* as soluble recombinant proteins and used for mouse immunization.

| CDS <sup>1</sup> | Product <sup>2</sup> | AA region cloned in *E. coli* <sup>3</sup> | Size (AA) <sup>4</sup> |
| --- | --- | --- | --- |
| MBOVPG45_0019 | Carbohydrate uptake ABC transporter-2 (CUT2) family, permease protein | 365 - 662 | 298 |
| MBOVPG45_0117 | Putative lipoprotein | 26 - 326 | 301 |
| MBOVPG45_0311 | Membrane lipoprotein P81 | 27 - 728 | 702 |
| MBOVPG45_0370 | Putative membrane protein | 108 - 350 | 243 |
| MBOVPG45_0411 | Translation elongation factor Tu | 5 - 396 | 392 |
| MBOVPG45_0584 | Putative lipoprotein | 85 - 355 | 271 |
| MBOVPG45_0643 | Putative lipoprotein | 25 - 232 | 208 |
| MBOVPG45_0825 | Putative lipoprotein | 28 - 383 | 356 |

<sup>1</sup> Mnemonic of CDS in *M. bovis* strain PG45 (GenBank: CP002188.1). <sup>2</sup> Protein products in GenBank CP002188.1. <sup>3</sup> The corresponding amino acid (AA) region was expressed in *E. coli* as a maltose-binding protein (MBP) fusion protein using the pMAL<sup>TM</sup>-c5X expression vector (New England Biolabs). When necessary, mycoplasma UGA tryptophan codons were converted to UGG codons to prevent premature translation termination in *E. coli*. <sup>4</sup> The size of the PG45 AA region cloned in *E. coli* is indicated.

Male Holstein calves (3 months old) originating from southwestern France (Tarn-et-Garonne) were housed at ENVT animal facilities. Serological analyses were performed using in-house ELISA (see below). At week 21, animals were immunized by a single subcutaneous injection of a heat-inactivated preparation of *M. bovis* PG45 (2 × 10⁹ CFU equivalents per animal) emulsified in Montanide ISA 61 (Seppic). Sera were collected at regular intervals from week 4 to 26. Normal Bovine Serum was purchased from a commercial vendor (Sigma, reference number B8655, Lot 127H9001) and is described as “a pooled preparation obtained from a normal donor herd”. Normal Mouse serum was purchased from a commercial vendor (Sigma, reference number M5905, Lot 0000480419). The anti-Strep monoclonal antibody was purchased from a commercial vendor (IBA Lifesciences; reference number 2-1507-001, lot 15707003) and is described as “a murine monoclonal antibody (IgG1 subclass) and highly selective for Strep-tag II tagged proteins”. The anti-HA monoclonal antibody was purchased from a commercial vendor (Invitrogen; catalog number 26183; lot ZF393513) and is described as “a mouse monoclonal antibody, clone 2-2.2.14 (IgG1 isotype), which is specific for the HA peptide”.

### Serological analyses by ELISA

For ELISA-based screening of antibodies present in bovine sera, 96-well microplates were coated overnight at 4 °C with 5 × 10⁷ CFU of *M. bovis* PG45 per well diluted in 0.05 M carbonate–bicarbonate buffer (pH 9.6). Plates were washed five times with DPBS containing 0.05% Tween 20 (PBST) and blocked with 300 µL of 5% non-fat dried milk (Sigma) in PBST for 2 h at 37 °C. Plates were then incubated for 1 h at 37 °C with serial dilutions of sera starting at 1:20. After five washes with PBST, horseradish peroxidase-conjugated goat anti-bovine IgG (Abcam), diluted 1:2,000, was added and incubated for 1 h at 37 °C. Following five additional washes with PBST, 3,3′,5,5′-tetramethylbenzidine (TMB) substrate was added and incubated for 10 min at 37 °C. The reaction was stopped by adding 50 µL of 2 M sulfuric acid per well, and absorbance was measured at 450 nm using a microplate reader (Bio-Rad, Hercules, USA).

## Results

### The BoviScan library design covers the majority of the *M. bovis* pan-proteome

To design the BoviScan library, we first selected a set of representative *M. bovis* genomes available either through the NCBI Genomes database or through Molligen^49^. The protein coding sequences from 566 genomes representing 295 strains were collected (Table S1).

This dataset covers a large section of the genetic diversity of *M. bovis* available worldwide (Figure 2A). It is comprised predominantly of strains isolated from cows (66%) and bison (32%) but also includes a small set of strains originating from atypical hosts such as deer or caprines (2%). Annotation for the tissue of origin was available for most strains (78%), with as expected a predominance of respiratory (52%) and mammary (15%) isolates, as well as a smaller set of less typical isolates from internal organs, joints, and skin and ears. Geographically, these isolates originated mostly from North America (64%), China (18%) and Europe (14%).

**Figure 2:**
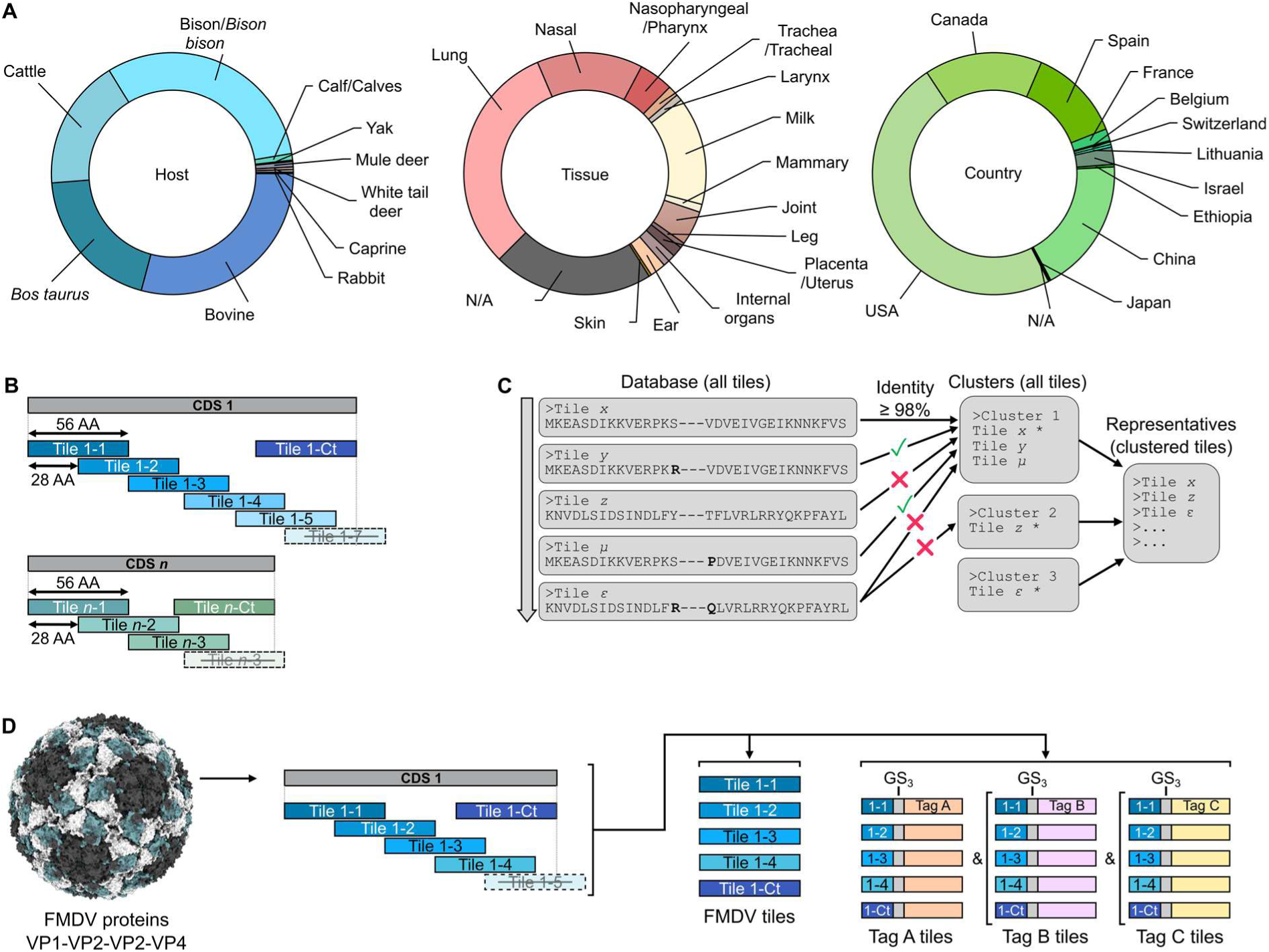
Composition of the BoviScan library. A. A set of genomes originating from 295 *M. bovis* strains were used a source of protein coding sequences to design BoviScan. For each strain the host, tissue and country of isolation were reported, and are summarized in the corresponding doughnut charts. N/A: not available. B. Schematic representation of the *pepsyn* tiling process. Each coding sequence is split in 56 amino-acids long tiles, overlapping each other by 28 amino-acids. The final tile covering the C-terminal end of the coding sequence often does not match the required 56 amino-acids lengths, and is therefore discarded. Instead, a specific C-terminal tile, starting from the Stop codon is generated. C. Overview of the CD-HIT clustering logic, when all sequences to cluster are the same length. Sequences are processed in their order of appearance in the input fasta file. By default, the first sequence is considered to be the representative of the first cluster. Any sequence matching the first cluster representative to the required identity threshold is clustered with it. The first sequence below the required threshold becomes the representative of the second cluster. Eventually, all sequences are part of a cluster, and a final list of cluster representatives is generated as output. D. A set of control peptides was generated using the coding sequences for the four structural proteins of the Foot-and-Mouth Disease virus. In addition, each of the FMDV peptide was appended with an epitope tag, fused via a short (GS)_3_ linker, and trimmed from the N-terminal side to match the 56 amino-acids length.

The resulting dataset is comprised of 4,227,733 CDSs. Each CDS was divided into small, 56 amino-acid long, overlapping peptides called “tiles” compatible with phage display (Figure 2B). Specific C-terminal tiles covering the end of each CDS were added, to avoid shorter peptides. Using this logic, 5,140,114 tiles were generated, corresponding to 4,718,311 ORF tiles and 421,803 C-terminal tiles.

Given the high degree of sequence conservation across strains, a significant portion of the tiles generated are either duplicates or very similar. To eliminate this redundancy, the clustering algorithm CD-HIT^47^ was used to bin tiles together based on a sequence identity percentage (Figure 2C). We elected to cluster the tiles at 98% identity, which corresponds to a maximum of 1 amino-acid mismatch over a 56 amino-acids sequence. As all the sequences to cluster were the same length, the order in which they were clustered is relevant. Indeed, under this condition, CD-HIT will automatically use the first sequence processed as the first cluster representative sequence. Subsequently, each following sequence will be compared to the representative sequences found before it, and sorted as either redundant or a new representative. We therefore elected to start the clustering process with the tiles obtained from the reference genome of *M. bovis* type strain PG45 (GCA_000183385). This ensured that most of the tiles remaining after clustering originate from the *M. bovis* type strain.

The clustering process yielded a total of 37,027 tiles, comprised of 31,723 ORF tiles and 5,304 C-terminal tiles. This corresponds to a 99.27% reduction compared to the non-clustered tiles set. This level of compaction was expected, given the high degree of genetic similarity observed between different *M. bovis* strains. We note that the library post-clustering contains tiles from only 478 genomes and 291 strains (Table S1). This is due to the presence in the initial dataset of two sets of genome annotations (RefSeq and GenBank submitter) for most strains. If both annotations for a strain had no difference, then one of them was fully removed by the clustering. We also note that clustering had no significant effect on the strain’s diversity representation. Indeed, at least one tile from most strains was conserved in the post-clustering library. Therefore, the host, tissue or country of origin distributions were similar pre- and post-clustering (Figure S1).

Finally, in addition to the tiles from *M. bovis*, the BoviScan library design also includes a set of tiles to be used as potential known-negative and/or known-positive controls (Figure 2D). The first set corresponds to peptides from the four proteins forming the viral particle of the Foot-and-Mouth Disease Virus. The second set corresponds to the FMDV tiles fused in C-Terminal to a well-defined epitope tag (HA tag, c-Myc tag, Avi-tag, StrepTag ii, and ALFA tag) *via* a (GS)_3_ linker. In order to maintain the 56 amino-acids length, the FMDV portion of the tagged-tiles was trimmed down from the N-terminal side. A total of 43 FMDV tiles and 215 FMDV-Tag tiles were generated and appended to the BoviScan design.

The final design of BoviScan is thus comprised of 37,285 tiles.

### Cloning and quality control of the BoviScan library

The amino-acid sequences of the 37,285 tiles were reverse-translated into DNA coding sequences. Each tile coding sequence (168 nucleotides) was appended with both a prefix and suffix sequences in 5’ and 3’, respectively (Figure 3A). The resulting set of 200 nt sequences was synthesized as an oligonucleotides pool and converted to dsDNA by PCR, using primers matching the prefix and suffix sequences. After amplification, a sample of the library was sequenced in order verify that the DNA pool was matching the BoviScan library design and that each coding sequence was present in similar amount. After stringent mapping of the sequencing reads the reference library, we observed that only 8 of the 37,285 sequences were not found in the data (Table S2), indicating that 99.98% of the expected library was present. Sequence abundances were also satisfactory, with 96.24% of the expected sequences found within 1 log of abundance (Figure 3B).

**Figure 3:**
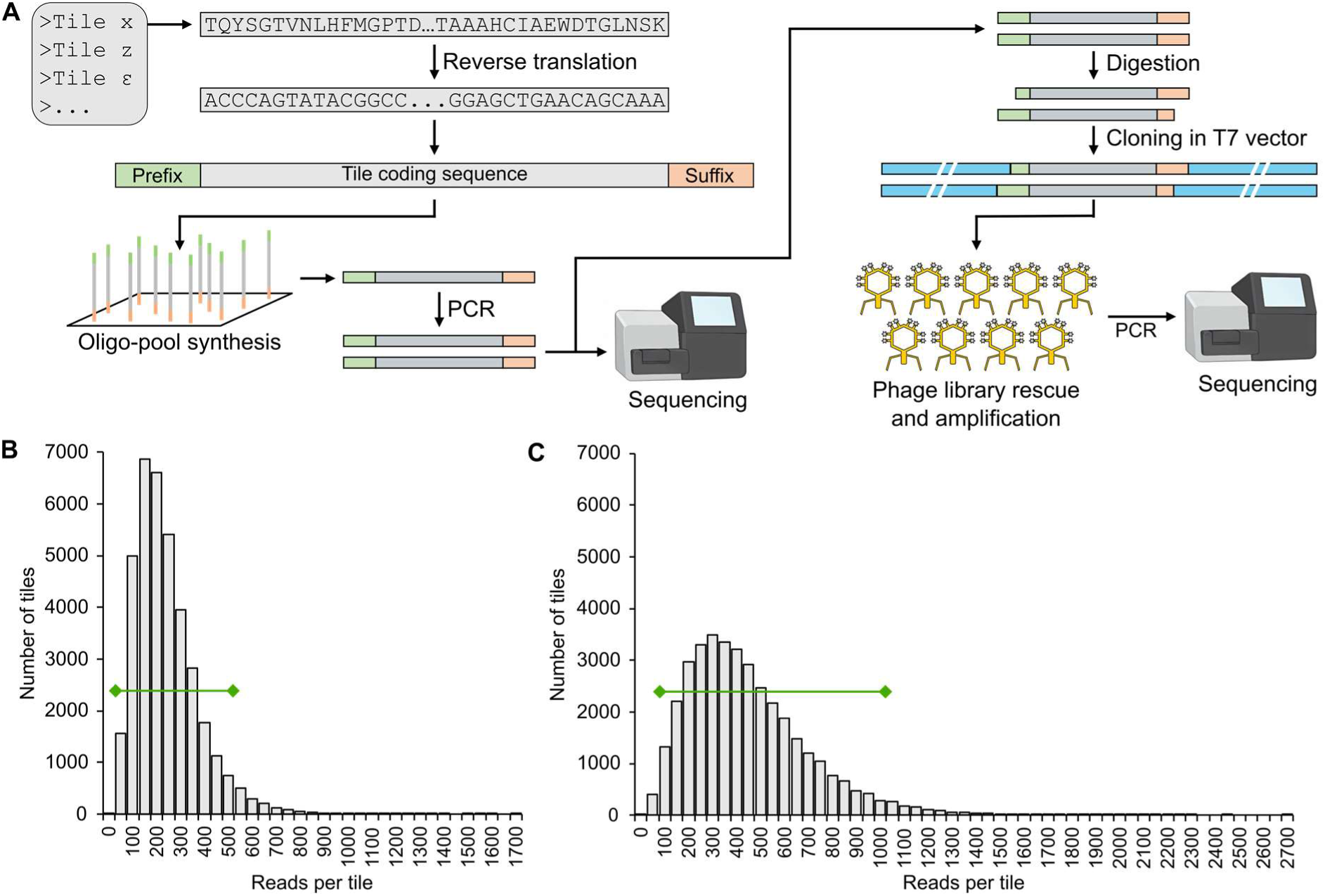
Quality control sequencing of the BoviScan library. A. Following the peptide clustering step, the representative sequences are reverse translated into coding sequences, append with identical 5’ and 3’ extremities (Prefix and Suffix), and synthetised as an oligonucleotides pool. The single stranded DNA is amplified by PCR, and sequenced to check that the synthesis matches the expected design. The DNA is then digestion by restriction enzymes, and ligated into the T7 bacteriophage genome to create a phage display library. The content of the phage library is checked by sequencing. B. Distribution of the number of reads mapped to each tile in the amplified oligo-pool sequencing data. Green line: 95% percent of the distribution. C. Distribution of the number of reads mapped to each tile in the cloned library sequencing data. Green line: 95% percent of the distribution.

The DNA library was then cloned into T7 viral particles, which were subsequently rescued and amplified in *E. coli*, yielding the final BoviScan phage library titrated at 7.5x10^10^ pfu.mL^-1^. To check that the cloning process did not negatively skew the library composition and distribution, sequencing was performed again. The tile-encoding locus in the phage genome was amplified by PCR, and the amplicon were sequenced and mapped to the reference design. Overall, only 12 tiles were not found in the dataset (Table S3), including 4 that were absent from the pre-cloning results. This corresponds to a 99.97% coverage of the BoviScan design. Abundance distribution was also correct, with 95.69% of the sequences found within 1 log of abundance, although the distribution was noticeably wider than pre-cloning (Figure 3C). In addition, we observed that ∼79.8% of the sequencing reads were perfectly aligned to the reference, indicating that most of the library was cloned in-frame and without mismatch or indel. This was confirmed by performing Sanger sequencing of the region of interest in a set of 24 plaque forming units. Two clones presented each a single missense mutation in the tile coding sequence, one clone presented a concatenated tile coding sequence (probably due to improper ligation), whereas the remaining 21 clones (87.5%) were identical to the library design.

Given these results, the final phage library was properly cloned, matching the BoviScan library design, and was thus used for subsequent immunoprecipitation experiments.

### Mock-immunoprecipitations do not yield any specific or spurious peptide enrichment

During PhIP-Seq experiments, a number of samples are dedicated to controls called “mock-immunoprecipitation” (mock-IP). In these mock-IPs, immunoglobulin-containing samples are omitted in order to assess whether any tile from the library is inappropriately enriched during the immune-precipitation process. These mock-IP samples also provide a background noise information, enabling by comparison the quantification of the antibody-specific enrichment in regular samples. This background is linked to the non-specific binding of the phages to magnetic beads used for the immunoprecipitation^29^.

Two sets of duplicate mock-IPs were performed independently. In all mock-IPs, reads from the vast majority of the tiles were recovered (99.44-99.91% coverage) (Table S4). The number of reads mapped for each tile in the four samples had a similar distribution to that of the phage library, and no tile was over-represented. Pairwise comparisons indicated that the mock-IP were reproducible, with good correlation between the assays (linear regression coefficient: 0.928-0.968, R^2^: 0.879-0.938; Pearson *r*: 0.803-0898) (Figure S2).

These results validated the immune-precipitation process, as it did not yield any unwarranted signal and no specific interaction occurred in these conditions between the phages library and the magnetic beads used. Thus, any enrichment observed for antibody-containing samples should be linked to specific interactions with the epitopes.

### Technical replicates of BoviScan PhIP-Seq experiments yield similar results

In order to assess the technical robustness of our experimental workflow, two sets of duplicate samples were processed across two independent experiments, analysed and compared. In these assays, a single aliquot of “normal bovine serum” purchased from a commercial vendor was used as sample.

Compared to the mock-IPs, the numbers of mapped reads for each tile followed a distinct distribution, marked by a small set of tiles with very high read counts (Figure S3A and Table S4). The maximum read count were an order of magnitude higher (5966.8-9533.5, compared to 445.5-651.1 in mock-IPs) and the number of tiles with read count 10-fold above the median was also vastly higher (241-289 tiles, compared to 0-1 tiles in mock-IPs). These observations are expected from PhIP-Seq samples in which peptide enrichment has occurred. In this case, it is highly probable that the commercial bovine serum used contained either cross-reactive anti-*M. bovis* antibodies or true anti-*M. bovis* antibodies resulting from infection of one or more animals from the source herd.

Pairwise comparison of the numbers of reads mapped for each tile showed that results obtained in the four assays were very similar, with a strong correlation between replicates (linear regression coefficient: 0.909-1.16, R^2^: 0.902-0.967; Pearson *r*: 0.937-0.981) (Figure S3B), indicating that similar enrichment was occurring in all samples.

Given this high reproducibility, and in the frame of this pilot study, we elected to perform the remainder of our experiments with a single PhIP-Seq assay per sample to analyse.

### Benchmarking BoviScan using monoclonal antibodies and protein-specific polyclonal antibodies

To assess the type of data that can be obtained with BoviScan, we elected to first analyse a set of known-positive samples containing antibodies directed against specific epitopes or specific protein antigens. The first two samples consist of monoclonal mouse immunoglobulins targeting the StrepTag and the HA-tag peptides, respectively. The third sample consists of a mix of sera collected from mice immunized with one of eight different recombinant proteins derived from *M. bovis* PG45 (Table 1). With these 3 samples, and in contrast with the mock-IPs, we observed that a small number of tiles (n=32-55) had extremely high numbers of mapped reads (normalized RC >1000, for a median normalized RC = 44) (Figure S4 and Table S4). These results are the expected outcome of antibody-mediated tile enrichment. To further quantify the peptide enrichment, the number of reads mapped for each tile in the samples was compared to the values obtained in the mock-IPs. To do so, EdgeR was used to generate a list of statistically enriched tiles (Table S5). In addition, a Z-score was calculated for each tile^50^, based on the mean and standard deviation of read counts for this tile in the mock-IPs (Table S6). A null Z-score denotes a value unchanged from the mock-IP, while a positive Z-score indicates an enrichment and negative Z-score indicates a depletion. The higher the Z-score absolute value, the higher the deviation from the mock-IP reference values.

In the Figure 4, both the read counts, and EdgeR and Z-score enrichment are displayed for each tile. In this data representation, the tiles are organized along the x-axis by their “Tile index”. This index is attributed based on the functional annotation of each tile, using on the reference proteome of *M. bovis* PG45, the relative position of each tile in the corresponding protein, and the position of each protein in the genome. Using this approach, tiles are ordered in a manner that indicates their positions in the genome of *M. bovis*, with “Tile index 1” corresponding to amino-acids 1-56 of the protein MBOVPG45_0001, “Tile index 2” to the tile covering the amino-acids 28-84; and so on. Using this approach, tiles indexed 1-32372 correspond to those that are similar to proteins MBOVPG45_0001-MBOVPG45_0902. Meanwhile, tiles indexed 32372-37027 correspond to those with no similarity to *M. bovis* PG45 proteome; and tiles indexed 37028-37285 correspond to Control tiles.

**Figure 4:**
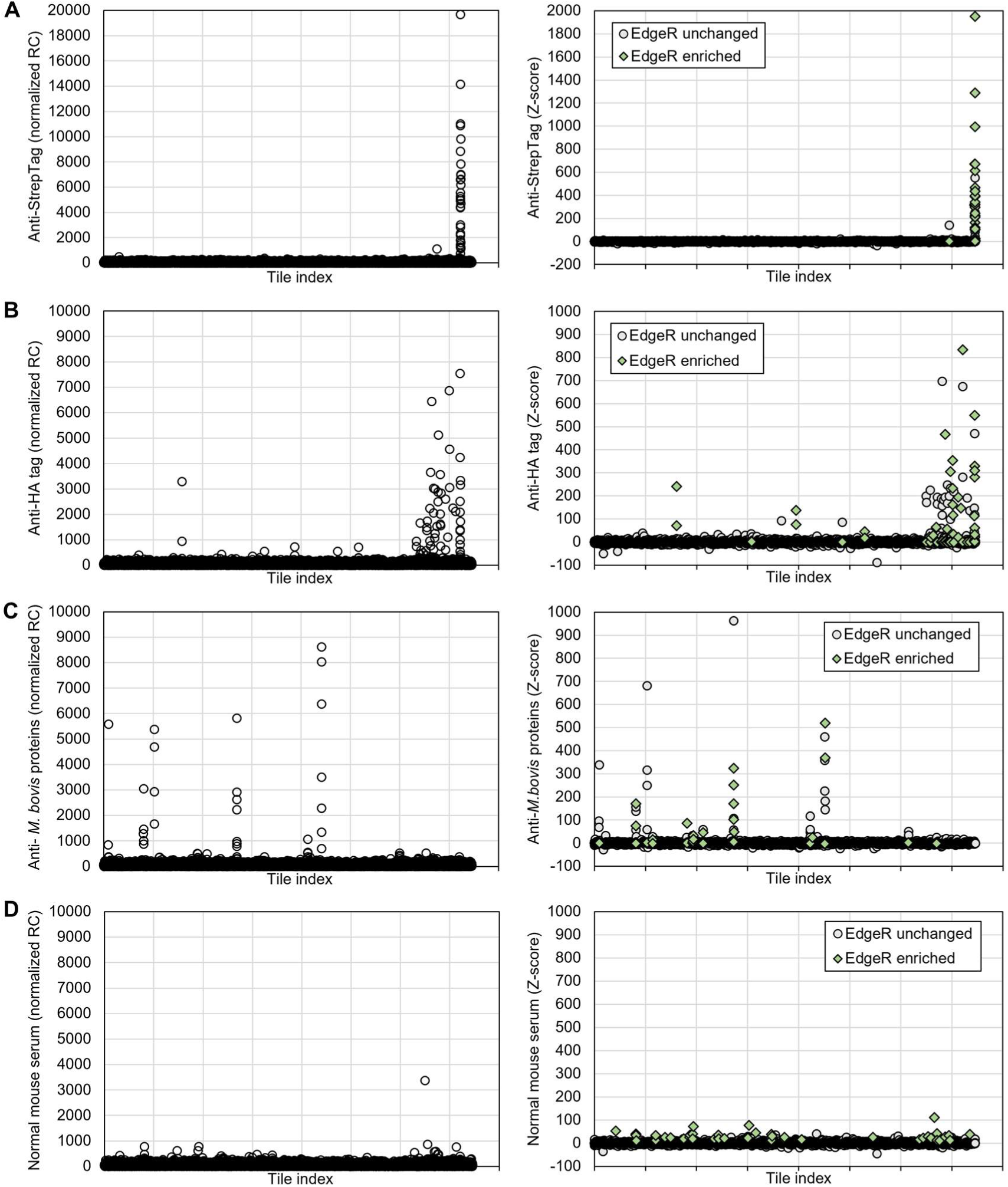
Peptide enrichment in BoviScan using mouse antibodies targeting known antigens. PhIP-Seq was performed using mouse mono-clonal antibodies, and sera from mice immunized with recombinant *M. bovis* proteins. The number of sequencing reads mapped to each peptide coding tile is presented (left). The tiles are organised in a specific order that mimic the repartition of the corresponding proteins along the genome of *M. bovis* PG45. In this arrangement, the first tile corresponds to the first 56 amino-acids of the protein DnaA. Tiles #1-32372 corresponds to proteins MBOVPG45_0001 to MBOVPG45_0902. Tiles #32373-37027 correspond to peptides with no matches in *M. bovis* PG45. Tiles #37028-37285 correspond to control tiles. In addition, the calculated Z-score for each tile is provided (right). Tiles annotated as enriched by EdgeR are displayed as green diamonds. A. Peptide enrichment in BoviScan using a mouse monoclonal IgG anti-StreTag II. B. Peptide enrichment in BoviScan using a mouse monoclonal IgG anti-HA tag. C. Peptide enrichment in BoviScan using a mix of 8 sera from mice, each immunized with a different recombinant *M. bovis* protein (Table 1). D. Peptide enrichment in BoviScan using normal mouse serum.

In the anti-StrepTag sample (Figure 4A), 34 tiles were considered significantly enriched by EdgeR, and 52 tiles had a high calculated Z-score (Z-score >10, for a median Z-score of -0.37). The majority of tiles identified as enriched corresponded to StrepTag tiles (33/34 EdgeR hits and 33/52 high Z-score), and no other control tile was considered enriched. All the StrepTag control tiles (43/43) were recovered in the sample, and the majority was considered enriched (33/43 by EdgeR; 33/43 by Z-score). However, some StrepTag tiles were not enriched, despite being present. This may be due to the fact that each of the 43 tiles has a distinct N-terminal sequence, which may play a role in the peptide folding and accessibility, thus potentially masking some of the epitope tags.

For the anti-HA Tag sample (Figure 4B), the results were less conclusive. While all the HA-Tag control tiles were recovered in the sample (43/43) they only accounted for a small fraction of the EdgeR enriched tiles (15/77) or high Z-score tiles (18/164 tiles). In addition, most HA-tag tiles were non-enriched, with only 18/43 tiles flagged by either method. While the same argument regarding the distinct N-terminal sequences impact on epitope masking could be applied, this enrichment rate was much lower than that observed for the StrepTag tiles. This could be also linked to epitope competition, as we observed that the majority of the enrichment corresponded to *M. bovis* tiles. Indeed, our data could be explained if the monoclonal anti-HA tag antibody had a relatively low specificity. In this case, as a portion of the antibodies would be bound to non-HA Tag tiles, there would be a corresponding reduction in signal for the HA Tag tiles. These results were confirmed in an independent replicate PhIP-Seq experiment (Figure S5), suggesting that the enrichment of *M. bovis* peptides by the monoclonal anti-HA antibody was repeatable.

For the anti-*M. bovis* proteins mixed sample (Figure 4C), the results obtained showed EdgeR enrichment and high Z-scores for 57 and 99 tiles respectively. As observed for the monoclonal antibodies, discrepancies existed between the two quantification methods, although a portion of the results overlapped (21 tiles). Interestingly, the majority of the tile enrichment was focused on a small set of only 10 *M. bovis* proteins (54/57 EdgeR enriched and 57/99 high Z-score). Enriched peptides matched 7 of the 8 antigens used to immunize the mice (MBOVPG45_0019, 0117, 0311, 0411, 0584, 0643, and 0825) and none was retrieved for one (MBOVPG45_0370). The other 3 proteins with tiles enrichment corresponded to housekeeping proteins (MBOVPG45_0158, 0293 and 0340). We also processed a sample of commercial normal mouse serum (Figure 4D) in order to check if some of the tile enrichment observed above was non-specific. While tile enrichment was detected (46 EdgeR hits and 165 high Z-score tiles), it did not match that of the immunized mice (no overlapping tiles in EdgeR and 10 overlapping tiles for high Z-scores). In addition, the Z-score values recorded where much lower (mean Z-score for tiles with Z-score >10: 20.2 for normal mouse serum and 84 for anti-*M. bovis* proteins serum), suggesting the absence of specific enrichment in this control sample.

Taken together, these benchmarking experiments suggest that the BoviScan library, coupled to our PhIP-Seq protocol, can provide a representation of the antigens targeted by antibodies found in a given sample. However, this representation can be skewed, depending in particular on the parameters used to assess tile enrichment. Here, we noted that while most of the high Z-score tiles were considered enriched by EdgeR, a portion was not; and conversely a number of low Z-score tiles were considered according to EdgeR. In addition, EdgeR annotated as “unchanged” a number of very high Z-score tiles, in particular in the anti-*M. bovis* proteins sample. We therefore elected to only use the Z-score as a measure of enrichment in the rest of the study.

To determine an empirical Z-score value threshold for enrichment, we performed a peptide enrichment analysis on the four normal bovine serum technical replicates (see above). The mean standard deviation was *σ* = 1.89, with a 95^th^ percentile of 5.09 and a 99^th^ percentile of 10.45. Meanwhile, analysis of pairwise differences showed a 95% confidence threshold of 7.10, and a 99% confidence threshold of 15.51. Based on these values, and on the Z-scores distribution observed in the anti-StrepTag and normal mouse serum samples, we elected to adopt a very conservative threshold of Z-score ≥ 20 for tile enrichment in the remainder of our pilot study. In subsequent analyses, non-enriched peptides were all attributed a Z-score of 1, in order to enable log transform of the enrichment data.

### Using BoviScan to compare the serum antibodies of multiple animals

To demonstrate the potential of BoviScan for comparing the immune response to *M. bovis* across different individuals, we analysed bovine hyperimmune sera collected from six calves naturally infected with *M. bovis* and subsequently immunized with strain PG45: five *M. bovis*-seronegative animals were exposed to a calf presenting otitis, a clinical sign commonly associated with *M. bovis* infection. At week 4, all animals were seronegative, except for the diseased calf (Figure 5A). By week 20, all animals had seroconverted and were seropositive. To further stimulate the immune response against *M. bovis*, the animals were immunized at week 21 by a single subcutaneous injection of a heat-inactivated preparation of *M. bovis* strain PG45. Serum samples collected at week 26 and confirmed to be seropositive by ELISA were subsequently processed by PhIP-Seq.

**Figure 5:**
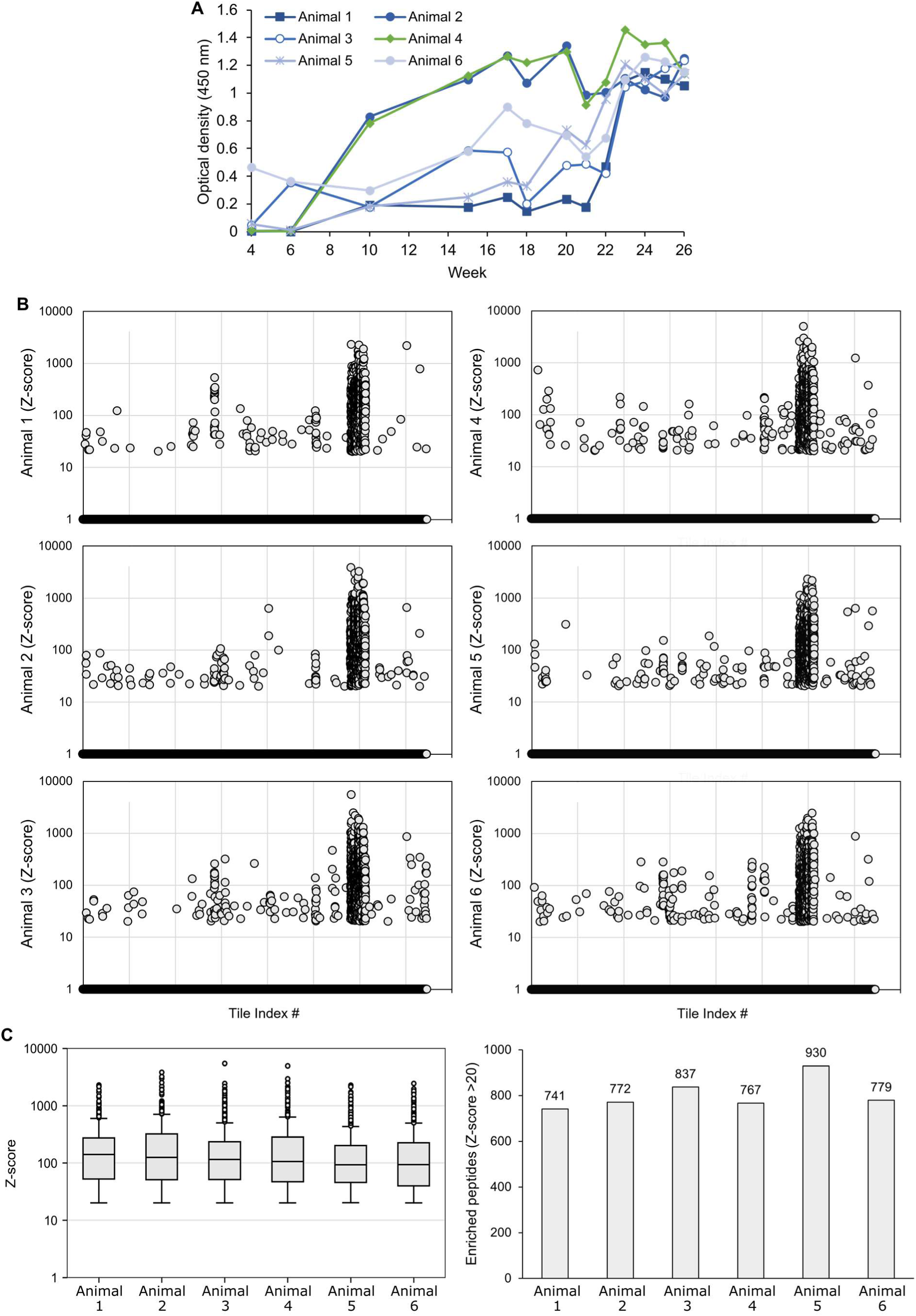
Comparative analysis of the sera from six experimentally immunized bovines using BoviScan. A. Serological analysis of the bovine serum samples by ELISA. For each of the 6 animals studied, samples were collected at various time points over 26 weeks and tested by ELISA for the presence of anti-*M. bovis* PG45 antibodies (IgG). B. Peptide enrichment in BoviScan using the sera from 6 animals collected at week 26. The tiles are organised in the same order as in Figure 4, matching the genome organisation of *M. bovis* PG45. For each tile-encoded peptide, the corresponding Z-score is reported. In order to enable log-scale display of the data, any peptide with a Z-score <20 (below the enrichment threshold) is attributed a Z-score =1. C. Box plot of the Z-score per peptide (left) and histogram of the number of enriched peptides (Z-score >20) in each of the 6 samples.

In all animals, a strong peptide enrichment could be observed (Figure 5B-C and Table S4, S6), with both a high number of enriched peptides (741-930 peptides with Z-score >20) and high Z-scores for these enriched peptides (median Z-score: ∼93-141; maximum Z-score: ∼2284-5456). Using the same Tile index representation as above, we observed that the vast majority of the enriched peptides originated from the same region, around Tile index 30000. These peptides share the functional annotation of “variable surface proteins”. In *M. bovis* PG45, 13 proteins form the *vsp* locus (MBOVPG45_0806-0813-MBOVPG45_0816-MBOVPG45_0818-0821). These peptides whose annotation matches any of these 13 proteins are hereafter termed “Vsp-like”. Depending on the sample, these Vsp-like peptides accounted for ∼80-87% of all enriched peptides (Figure S6). In addition, these were overall associated to higher Z-scores (median Z-scores Vsp peptides: ∼114-162; median Z-scores other peptides: ∼31-43).

Comparison of peptide enrichment patterns across the 6 animals revealed a high degree commonality (Figure 6A-B and Supplementary Figure S7), with 60% of the peptides shared by at least 2 animals and 30% shared by all animals. The Vsp-like annotated peptides represented the bulk of these common peptides, as they accounted for 98% of the 457 peptides common to all samples, 95% of the 96 peptides shared in 5 animals, 87% of the 110 peptides found in 4 animals, 76% of the 99 peptides common in 3 animals, and 62% of the 135 peptides shared in 2 animals. Conversely, peptides found only in a single animal, where predominantly derived from non Vsp-like proteins, and accounted for over 75% of the 597 unique peptides.

**Figure 6:**
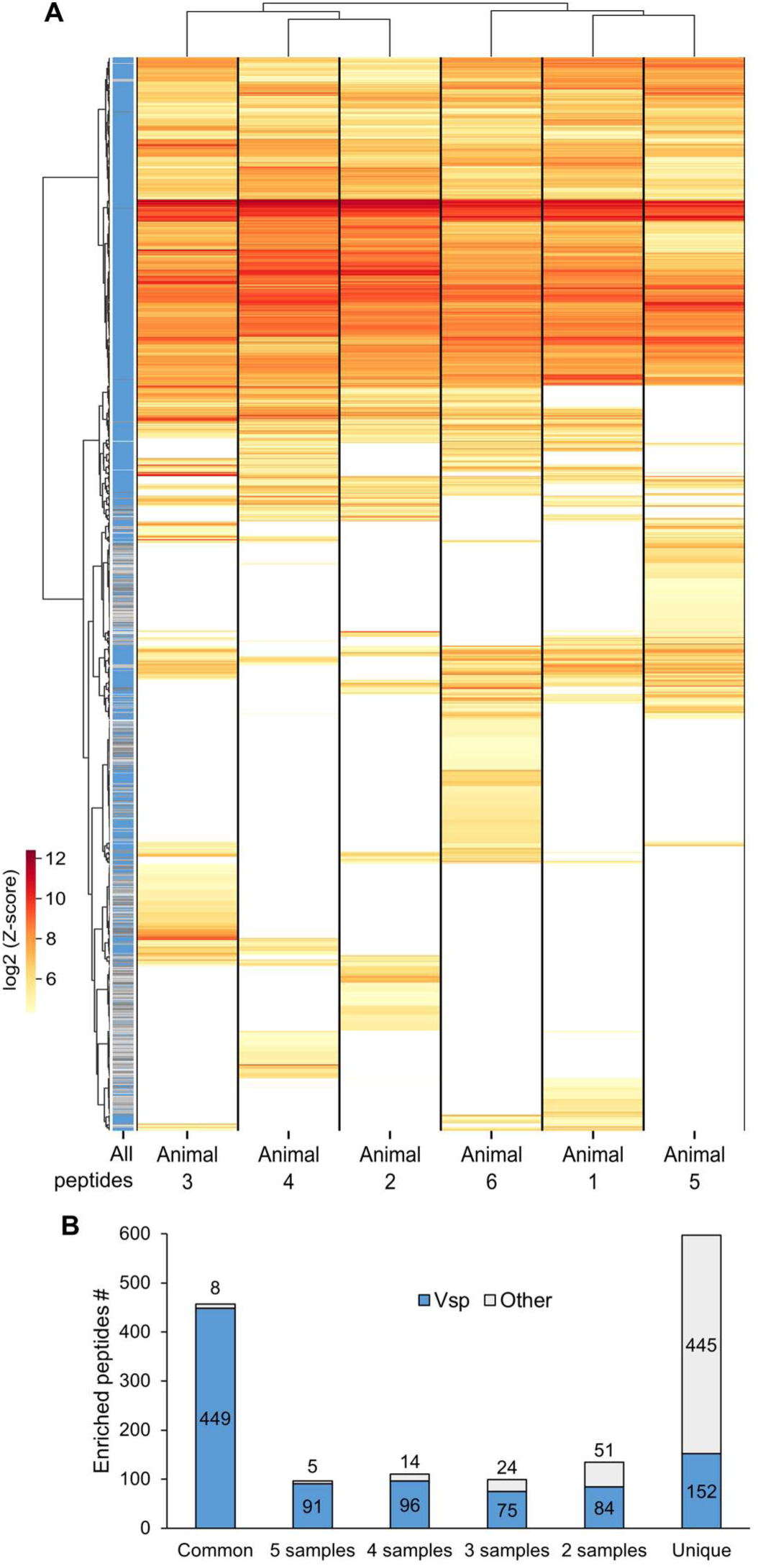
Comparison of peptide enrichment patterns across six animals using BoviScan. A. Heatmap of the enriched peptides in the 6 animal samples. The Z-score tables were filtered to keep only the peptides enriched in at least one sample. For any given peptide, Z-score values <20 were arbitrarily set to 1, to enable log conversion of the data. Hierarchical cluster analysis was performed on both the samples and peptides, using Ward’s method and Euclidian distance. The peptide functional annotation is indicated in the left-most column (blue: Vsp-like; grey: Other). B. Bar graph on the number of enriched peptides common to multiple animals, or unique to a single animal, split by peptide type (Vsp-like or Other).

Across the 6 samples, a total of 1494 peptides were considered enriched, with 947 of them (63%) corresponding to Vsp-like peptides. The other 547 enriched peptides, non Vsp-like annotated, carried annotations corresponding to a set of 149 proteins. Of these, 65 were “putative membrane protein” or “putative lipoprotein” and 9 were “conserved hypothetical protein”. Another 5 corresponded to the MIB-MIP virulence system, 6 to ABC transporters components, and 3 to the well-known surface proteins P48, P81 and MnuA. The remainder 60 proteins from the set corresponded to an array of cytoplasmic proteins involved predominantly in housekeeping, including subunits of the DNA polymerase III, the RNA polymerase, and various ribosomal proteins.

Interestingly, of these 149 protein annotations, only a small subset was the source of most the peptide enrichment and they were all surface associated. The most prevalent were MBOVPG45_0703 and MBOVPG45_0423, two putative lipoproteins, with corresponding peptides found highly enriched in all six animals. Next were MBOVPG45_0427 and MBOVPG45_0431, another pair of putative lipoproteins, with corresponding peptides found enriched in 4 and 5 of the animals, respectively. Peptides matching four other lipoproteins of unknown function were retrieved: MBOVPG45_0038 (PDxFFG protein), MBOVPG45_0679 (putative membrane protein), MBOVPG45_0509 (BspA family leucine-rich repeat surface protein) and MBOVPG45_0481 (putative membrane protein ICEB-1 encoded). Enriched peptides matching two well characterized virulence factors were also found: MBOVPG45_0376, a part of the MIB-MIP system; and MBOVPG45_0710, the lipase MilA. Both were retrieved in multiple animals (5/6 and 4/6, respectively) albeit with relatively low peptide counts (1-6 and 1-5, respectively) and low-to-medium cumulated Z-scores (25-256 and 27-242, respectively).

### Using BoviScan to perform a time-course analysis of the immune response

Animals used to produce hyperimmune sera were monitored over a 26-week period and regularly sampled for ELISA testing. Animal 2 was selected for longitudinal analysis of the immune response, since this animal was initially seronegative (week 4 and 6), before seroconverting and presenting high titres of anti-*M. bovis* antibodies (Figure 5A). Serum samples collected at weeks 4, 6, 10, 15, 17, 20, 23, and 25 were analysed using BoviScan (Table S4, S6). Our data showed that at week 4 and 6, while the ELISA were reported as negative (below threshold), a small set of peptides where enriched albeit at relatively low Z-scores (Figure 7A). Then, at week 10, the number of enriched peptides increased sharply, as well as the associated Z-scores. These values kept increasing over the following weeks, eventually reaching a maximum at week 17. This pattern matched that of the ELISA, and correspond to the immune response associate to the natural infection of the animal. At week 20, the overall number of enriched peptides was strongly decreased, together with the cumulated Z-scores, reverting to levels similar to week 10. Then, following immunization at week 21, these values increased again at week 23 and 25. This, again, was similar to trend visible in the ELISA measurements.

**Figure 7:**
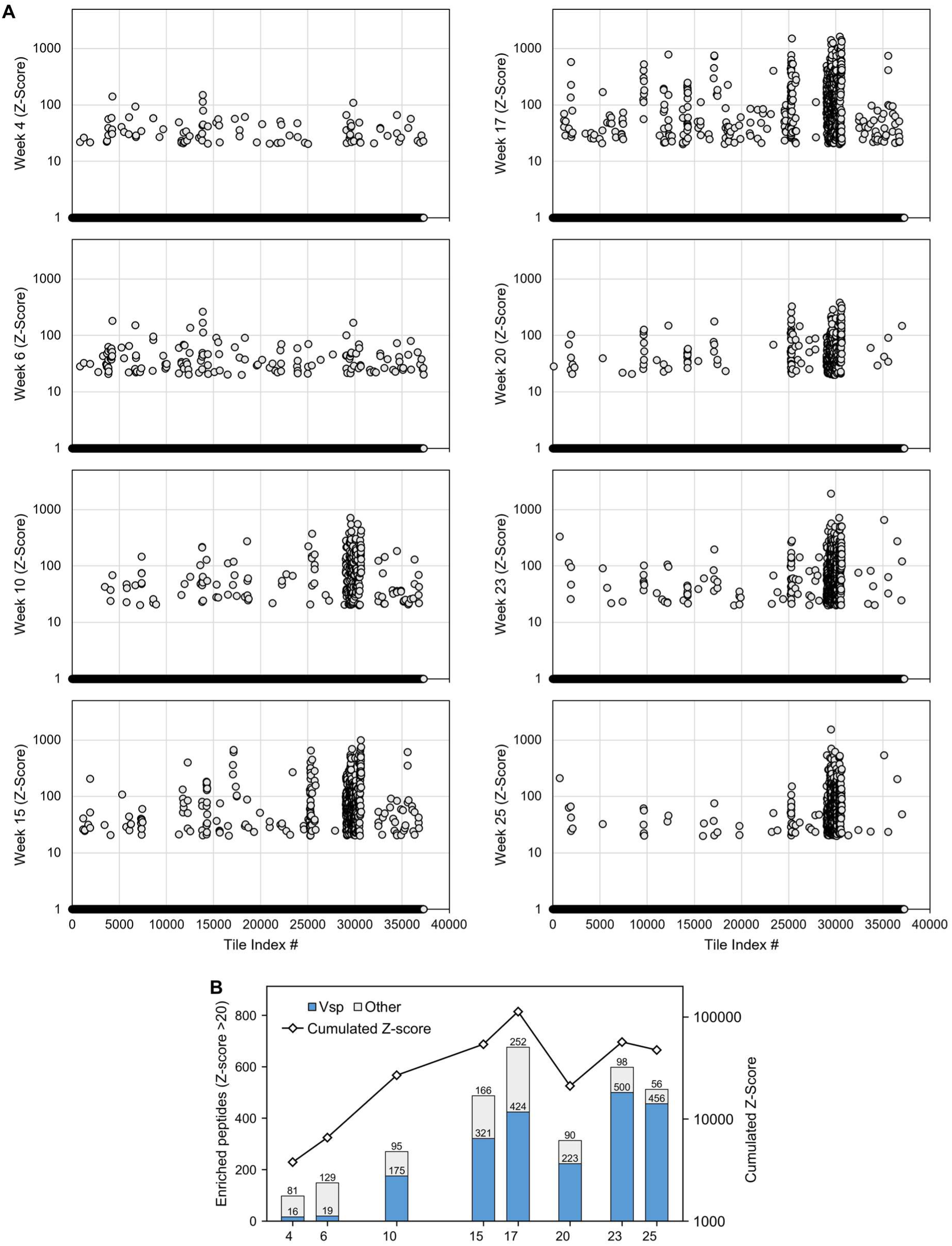
Comparative analysis of the sera from a single experimentally immunized bovine over 25 weeks using BoviScan. A. Peptide enrichment in BoviScan using the sera from one animal over a period of 25 weeks. The tiles are organised in the same order as Figure 4, matching the genome organisation of *M. bovis* PG45. For each tile-encoded peptide, the corresponding Z-score is reported. In order to enable log-scale display of the data, any peptide with a Z-score <20 (below the enrichment threshold) is attributed a Z-score =1. B. Left-axis: stacked bar-chart of the number of enriched peptides (Z-score >20) in each sample, split by peptide type (Vsp-like or other). Right-axis: line graph of the cumulated Z-score in each sample. The cumulated Z-score consist of the sum of the Z-scores for each enriched peptide in each sample.

Interestingly, this time series revealed that the high prevalence of Vsp-like annotated peptides observed at week 26 was not present for the initial phases of the immune response (Figure 7B). Indeed, the percentage of Vsp-like peptides was approximately 12-16% in weeks 4 and 6, and rose to a somewhat constant ∼62-65% in the later phases of the natural infection. However, following immunization, this percentage increased again, reaching over 80% in the last sample collected.

In terms of overall composition, it was also noticeable that the small set of peptides enriched during the early phases (week 4 and 6) was not found again in our dataset in the subsequent sampling points (Figure 8, Supplementary Figure S8). Similarly, a large part of the non Vsp-like peptides enrichment emerging at week 10-17 was then lost in the later stages of the longitudinal study. Finally, a specific set merged at week 23, dominated by a new pool Vsp-like annotated peptides.

**Figure 8:**
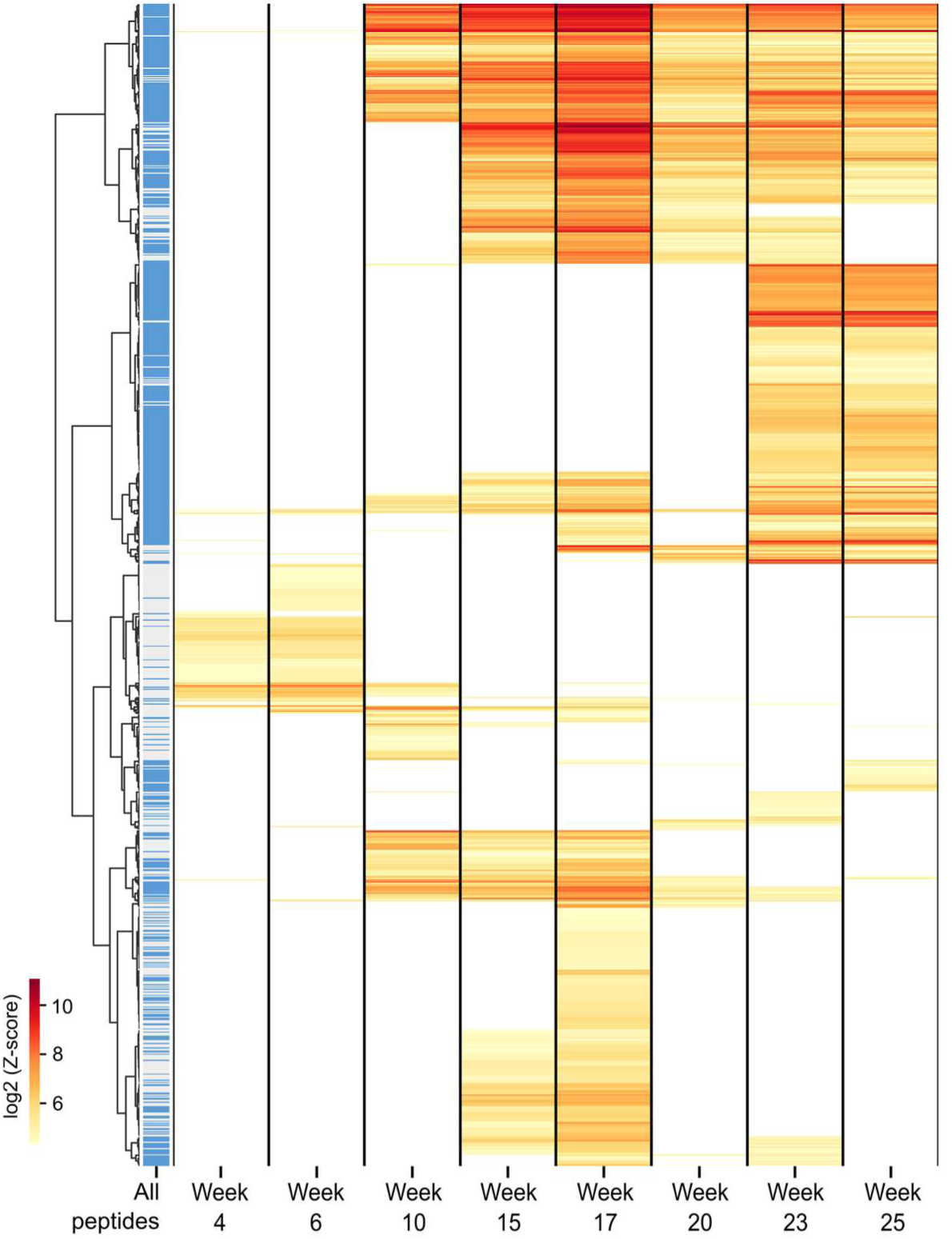
Comparison of peptide enrichment in a single animal during 25 weeks. Heatmap of the enriched peptides in the 8 samples of the time-series. The Z-score tables were filtered to keep only the peptides enriched in at least one sample. For any given peptide, Z-score values <20 were arbitrarily to 1, to enable log conversion of the data. Hierarchical cluster analysis was performed on both the samples and peptides, using Ward’s method and Euclidian distance. The peptide type is indicated in the left-most column (blue: Vsp-like; grey: Other).

Across the entire time series, enriched peptides with non Vsp-like annotations matching 127 distinct proteins were retrieved. However, this number was much lower at each time point (week 4: 36 proteins; week 6: 58 proteins; week 10: 31 proteins; week 15: 39 proteins; week 17: 53 proteins; week 20: 22 proteins; week 23: 22 proteins), and there was limited overlap between the different samples. This indicates that the composition of the antibody found in the animal serum was dynamic and varied strongly over the course of the experiment.

Some exceptions were noticeable. The first was MBOVPG45_0481 (a putative membrane protein, ICEB-1 encoded), for which enriched peptides with high Z-scores (mean Z-score: 158) were found at all time points. Then, a set of enriched peptides with annotations matching multiple putative lipoproteins (MBOVPG45_0038, MBOVPG45_0047, MBOVPG45_0159, MBOVPG45_0431), a MIP (MBOVPG45_0376) and the multifunctional lipase MilA (MBOVPG45_0710) were found enriched in 7 of the 8 time points analysed. Finally, peptides with annotations corresponding to the membrane lipoprotein P81 (MBOVPG45_0311) were found enriched both in the early (week 4-6) and late (week 17-25) phase samples, but not in weeks 10 and 15.

## Discussion

Our study was initiated with the goal of assessing whether PhIP-Seq could be employed as a complement to both ELISA and Western Blot, to better understand the antibody response developed by bovines during infection by *M. bovis*.

While multiple PhIP-Seq libraries are described in the literature, none was dedicated to the study of a given bacterial pathogen. This may be due in part to the fact that bacterial antigens often include lipopolysaccharides, in addition to membrane and secreted proteins. As PhIP-Seq relies on phage display of peptides, this technique is not able to map the antibody repertoire targeting non-protein antigens. For mycoplasmas, this is less problematic, given that these bacteria are wall-less and do not produce LPS. However, we do note that several mycoplasma species produce a polysaccharide capsule^51^, putatively involved in virulence and immune-evasion^51,52^. The production of antibodies targeting these capsular polysaccharides is currently not well characterized, although their immune-modulatory property has been highlighted^53,54^. Overall, the limitations of PhIP-Seq are well documented and understood^29,31^. These revolve predominantly around the constraints associated to the phage-display technology, and in particular to the short size of the peptides that can be encoded in oligonucleotides libraires and displayed on viral particles. As a result, PhIP-Seq can only capture the antibodies that target short, linear epitopes, thus losing the information for antibodies binding to larger and conformal epitopes. This method is also blind to post-translational modifications.

The design of the library (i.e., its peptide content) is also an important factor to consider, as once the library is synthesized and cloned, it cannot be edited. It is however possible to perform PhIP-Seq experiment by pooling together multiple libraries at the antibody-binding phase^55^. This approach has been successfully used to simultaneously probe both the human microbiota and allergens libraries. This strategy could therefore be used to enrich an initial library, by adjunction of a second library of differing composition. In our case, the BoviScan library covers 295 strains of *M. bovis*, based on the sequencing data available at the time of library design and synthesis. While this set of strains covers a large part of the genetic diversity of *M. bovis*, a large number of new genomes have been sequenced since (at the date of writing, assemblies for 835 strains were available in NCBI genome). It may thus be beneficial to analyse this newly available diversity and evaluate whether it is significant enough to warrant the creation of a second library. We also note that *M. bovis* infections often occurs in the context of multiple co-infections by both other viruses and bacteria, for instance in the case of the Bovine Respiratory Disease Complex (BRDC)^56^. It could therefore by biologically relevant to create a BRDCScan library, encompassing the proteome of all the different agents associated with this syndrome.

Another aspect to consider when using PhIP-Seq is the fact that the standard protocol relies on Protein G and/or Protein A-coupled magnetic beads to perform the immune-precipitation step. While these immunoglobulin-binding proteins (IBP) have good affinities for human IgG, their performances vary strongly for other species (including some veterinary relevant species) and immunoglobulin isotypes. IgM and secretory IgA are especially poorly captured by Protein G and Protein A^57,58^. This could be problematic for the study of *M. bovis* antibody response and that of other mycoplasmas spp. of veterinary importance, as these pathogens are mostly found in the mucosa where the bulk of the antibodies produced are IgM and secretory IgA^59^. In addition, having access to both IgM and IgG data could help better understand the seroconversion between both types during infection. In order to solve this issue caused by Protein G and Protein A, it is possible to add to the PhIP-Seq process an intermediate antibody (often an IgG “anti-target antibodies”), for which the IBP has high affinity. Alternatively, Protein A/G beads can be replaced by streptavidin-coated magnetic beads pre-loaded with isotype-specific, biotinylated capture antibodies^60^. PhIP-Seq is also compatible with a broad range of sample type, and is not limited to serum. It can be used to perform serological profiling on any immunoglobulin-containing samples, including broncho-alveolar lavages, saliva and milk^48^, the latter being particularly relevant for *M. bovis*^61^.

Finally, in comparison to ELISA, PhIP-Seq requires a longer and more complex experimental process. A typical experiment is performed over 2-3 days: one day for tube preparation and blocking, one day for serum-library binding and one day for PCR and amplicon purification. The sequencing library must then be prepared, and actual short read sequencing must be performed. We note that although complex, this process can be fully automated, as all steps are compatible with robot-assisted pipetting. Once sequencing data have been generated, their processing can be automated and usually requires a few minutes per sample for read mapping to the reference. As a result, PhIP-Seq is not intended as a diagnostics or point-of-care tool, but rather to be used to perform *post-hoc*, in-depth analysis of samples. In exchange for its limitations, PhIP-Seq offers the possibility to perform an in-depth characterization of the antibody repertoire at a very large scale, both in terms of number of samples and of number antigens.

In our case, we have showed that PhIP-Seq and the BoviScan library could be used to describe in a very detailed manner the IgG content against *M. bovis* in bovine serum samples. The immunoprecipitation protocol was both repeatable and free of artefact. Our data showed very good agreement between BoviScan output and classical ELISA, as the number of enriched peptides reported was proportional to the optical density. Concordance between ELISA and PhIP-Seq data has been demonstrated in previous studies for other libraries, and are a hallmark of the PhIP-Seq ability to properly quantify the antibody response^62,63^. In addition to this correlation, BoviScan produced an extremely detailed view of the anti-*M.bovis* antibody repertoire, highlighting its extremely dynamic nature, whereas ELISA could only provide an overview of sero-positivity. We also note that BoviScan yielded a significant number of enriched peptides even for sero-negative samples. PhIP-Seq is known for its high sensitivity, and thus offers an advantage over ELISA to explore the presence of low-abundance antibodies and map their target epitopes, especially in early stages of infection.

Overall, the data generated with BoviScan regarding which antigens were targeted appeared to be coherent, and in agreement with the literature. However, they should be considered cautiously. Indeed, we acknowledge that the bovine serum samples used in this study are not good representative of the typical *M. bovis* infection. Owing to their atypical nature, and the limited size of this animal cohort, the present study should not be used to draw any general conclusion regarding the immune response to *M. bovis*.

Variable surface lipoproteins represented the bulk of the targeted antigens, in particular those encoded in the *vsp* locus. This is in agreement with the literature, which showed that these lipoproteins are extremely abundant at the surface of *M. bovis* and are dominant antigens^11,64,65^. Interestingly, the biochemical nature of these Vsp proteins could also lead to an over-representation in PhIP-Seq data. Indeed, most Vsp are comprised of short repeated motives, measuring between 6 and 80 amino-acids. These small motives therefore can be for the most part fully represented in a single PhIP-Seq peptide (56 amino-acids). As a result, they are not subject to the same negative representation of conformal epitopes seen for larger proteins, and PhIP-Seq enrichment can be gathered for both immunoglobulins targeting linear epitopes and those targeting conformal epitopes. This over-representation could then have a knock-on effect of under-representing other enriched peptides, as fewer sequencing reads would be available to identify them in a given sample. Various strategies could be implemented to offset this phenomenon. First, higher number of reads could be collected per sample, coupled to specific filtering-out of Vsp matching reads. Alternatively, Vsp-specific depletion of the samples could be envisioned, by first incubating the sera with magnetic beads coupled to recombinant Vsp proteins or peptides.

In addition to the variable surface lipoproteins, our datasets also contained enriched peptides for multiple surface proteins. In particular, we note the lipase MilA (MBOVPG45_0710), which was found to be enriched across the majority of samples. This observation matches the fact that this specific protein has been reported as highly relevant for *M. bovis* serological diagnostics, as it was found to be frequently immunogenic and later used as the basis for a highly specific and sensitive ELISA assay^66,67^. We also note that at least two proteins, reported to be immune-dominants in the literature, are seldom found enriched in our datasets. For instance, enriched peptides corresponding to the lipoprotein P48 (MBOVPG45_0016)^68,69^ were found in 5 of the 6 animal samples, but low numbers of peptides (1-3) per samples, and medium Z-scores (40-125). Meanwhile, in the case of lipoprotein P81 (MBOVPG45_0311)^28,70^, only one animal showed significant enrichment of corresponding peptides (7 peptides, Z-scores: 31-216). Interestingly, this animal was the one used for the time-track analysis, which showed a much higher enrichment at week 17 (the same 7 peptides, with Z-scores 124-529) and a steady decrease of Z-scores over time. These observations suggest that BoviScan is able to detect antibodies targeting these antigens, although in the case of P48 a bias may cause under-representation. It is also possible that the immune response observed in the 6 animals at week 26 was skewed by the immunization at week 21, which may have substantially altered the antibody pool. A complete analysis of all animals across the whole experiment might therefore lead to a different representation of the expected antigens. We also note that conflicting data exist in the literature regarding the dominant antigenic nature of P48^71^, which could also explain its low abundance in our datasets.

Overall, based on the results gathered in this initial study, we propose that BoviScan represent an important and powerful tool for the study of the immune response to *M. bovis*. It should be used as a complement to ELISA, as the detailed view of antibody reactivity it provides could be highly valuable to further our understanding of this economically relevant pathogen. A BoviScan-based analysis of a large cohorts of naturally infected animals could help mapping the “normal” immunologic profile occurring during infection. Detailed characterization of targeted antigens in individuals exhibiting no or mild symptoms and comparison with individuals severely affected could lead to the identification of protective epitopes. Furthermore, BoviScan could enable a detailed analysis of immune reaction following vaccination, enabling a more comprehensive comparison of different vaccine formulation and administration route, eventually paving the way toward an efficient prevention and control strategy against *M. bovis*. In addition, PhIP-Seq could quickly be applied to other mycoplasmas, through the design and cloning of new library targeting veterinary and human health relevant species.

## Acknowledgement

We thank H. Benjamin Larman for his advice regarding the library design and the implementation of the PhIP-seq protocol. We are grateful to the scientific community working on the development and implementation of PhIP-Seq for publishing extremely detailed protocols, which were essential to the rapid and efficient adaptation of this technique on a new model species. The corresponding author is thankful to Dr. Hadid Hedrifing for fruitful discussion on immune-precipitation. This study was funded by the French Agence Nationale de la Recherche, under grant ANR-21-CE35-0008 RAMbo-V, and by the European Union’s Horizon 2020 Programme, under the Grant Agreement n°633184 SAPHIR.

## Material availability

All the reference sequences used for library design, as well as the final library are deposited online in the Zenodo repository (doi: 10.5281/zenodo.22749183). The BoviScan library will be provided freely to any interested party upon request, together with detailed protocols for library propagation, immune-precipitation and data-processing.

## Supplementary Figure

**Figure S1:**
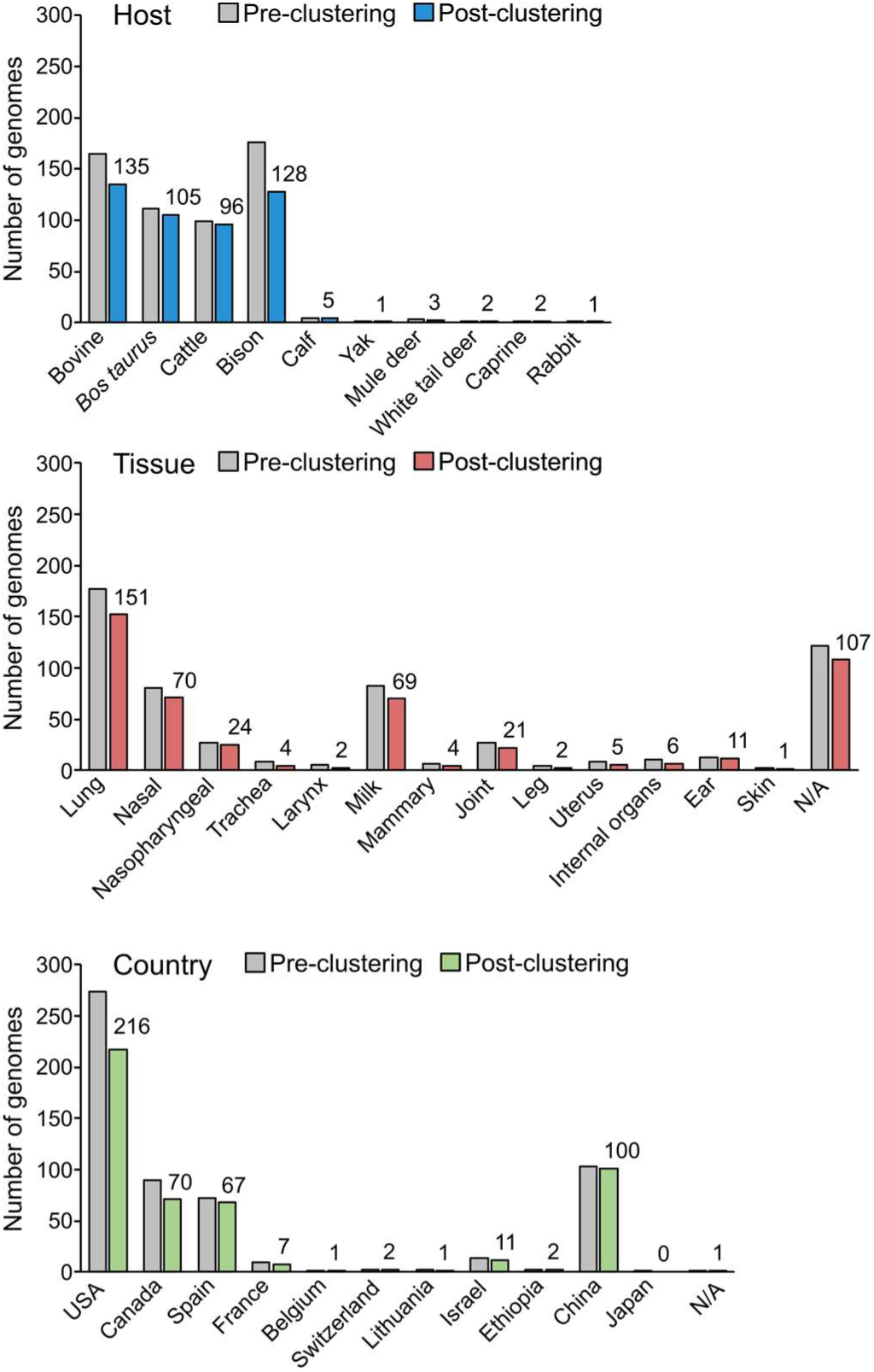
Comparison of the BoviScan library genome diversity before and after clustering of the peptides sequences. Bar-chart of the number of *M. bovis* genomes originating from a given host, tissue and country, in the Pre- and Post-clustering library. If at least one peptide originating from a given genome is present in the library, the corresponding source genome is accounted for. If no peptide from a given genome is found in the library, the corresponding source genome is not counted.

**Figure S2:**
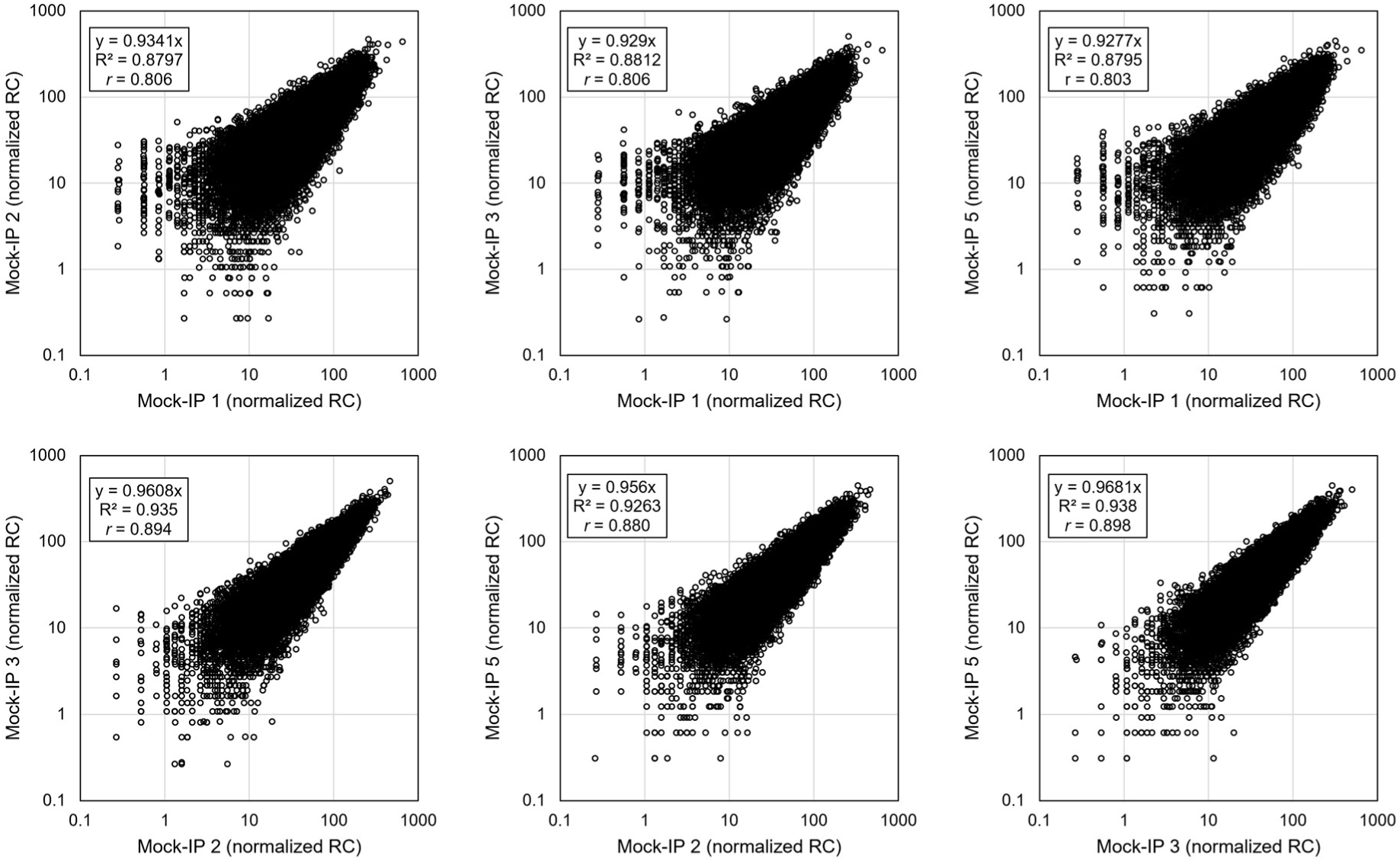
Comparison of the BoviScan data collected in the mock-IP samples. Pair-wise scatter plot of the normalized read count (RPM) obtained for each tile in the mock-IP samples. For each pair-wise comparison, linear regression coefficient (y), coefficient of determination (R^2^), and Pearson correlation coefficient (*r*) are indicated.

**Figure S3:**
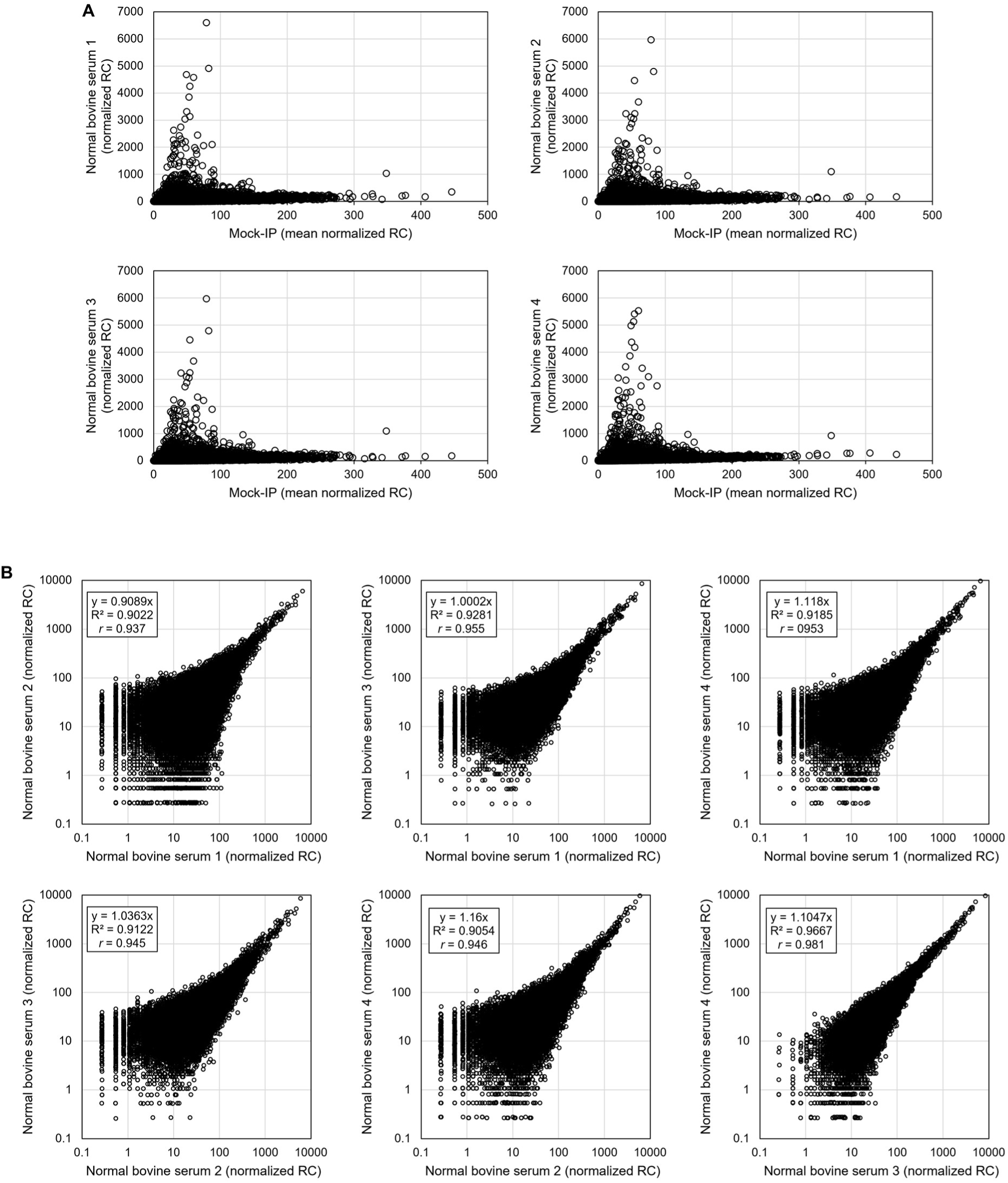
Comparison of the BoviScan data collected in four technical replicates analyses of normal bovine serum. A. Scatter plot of the normalized read count (RPM) obtained for each tile in the normal bovine serum samples and compared to the mean normalized read count (RPM) obtained in the mock-IPs samples. B. Pair-wise scatter plot of the normalized read count (RPM) obtained for each tile in the normal bovine serum samples. For each pair-wise comparison, linear regression coefficient (y), coefficient of determination (R^2^), and Pearson correlation coefficient (*r*) are indicated.

**Figure S4:**
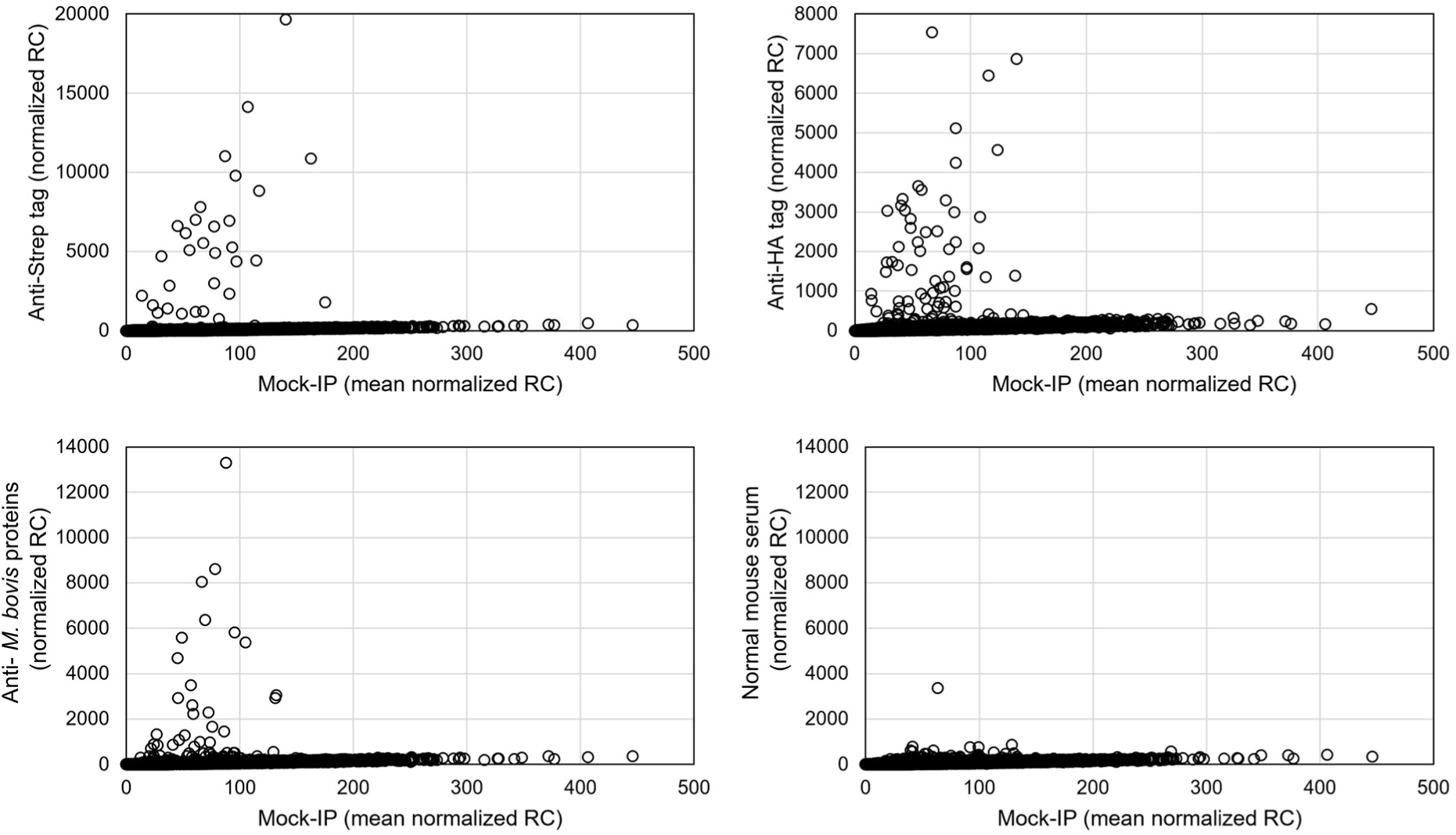
BoviScan data collected using mouse antibodies targeting known antigens. A. Scatter plot of the normalized read count (RPM) obtained for each tile in the Anti-Strep tag, Anti-Ha tag, Anti-*M. bovis* proteins and normal mouse serum samples and compared to the mean normalized read count (RPM) obtained in the mock-IPs samples.

**Figure S5:**
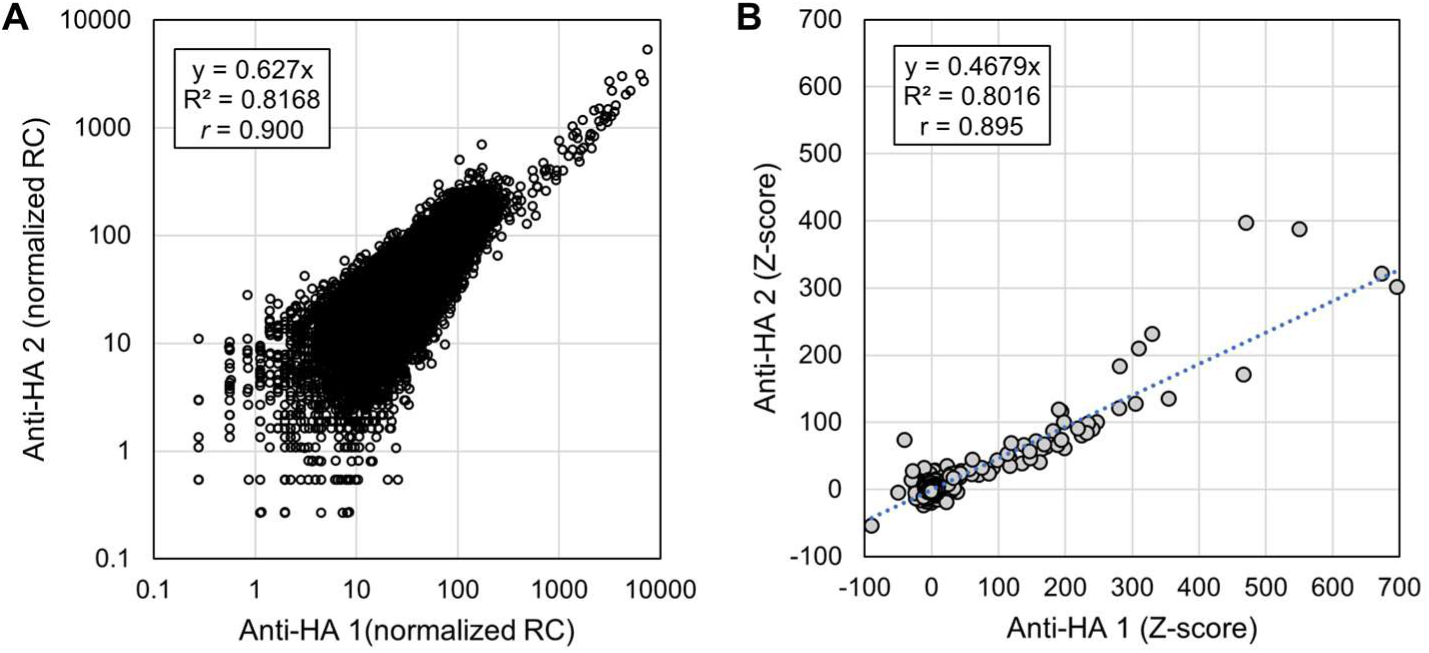
Comparison of the BoviScan data collected in two technical replicates analyses of the Anti-Ha antibodies. A. Scatter plot of the normalized read count (RPM) obtained for both replicates. Linear regression coefficient (y), coefficient of determination (R^2^), and Pearson correlation coefficient (*r*) are indicated. A. Scatter plotof the Z-score obtained for both replicates. Linear regression coefficient (y), coefficient of determination (R^2^), and Pearson correlation coefficient (*r*) are indicated.

**Figure S6:**
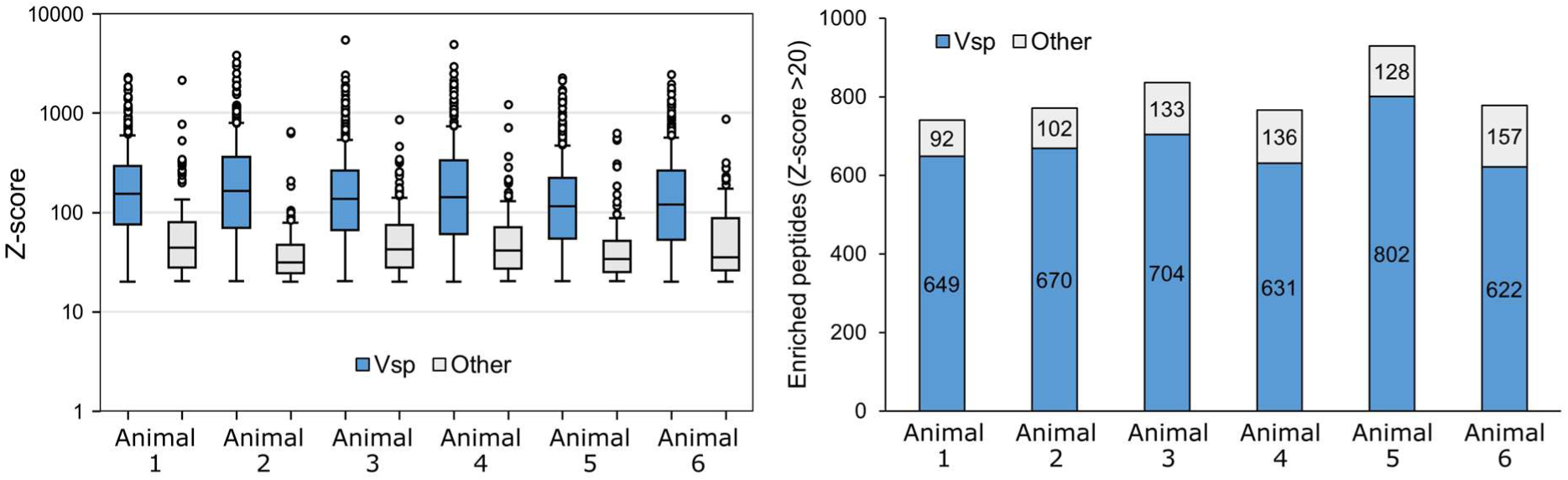
Comparative analysis of the sera from six experimentally immunized bovines using BoviScan. Box plot of the Z-score per peptide (left) and stacked bar-chart of the number of enriched peptides (Z-score >20), split by peptide type (Vsp-derived or other) in each of the 6 animal samples collected at week 26.

**Figure S7:**
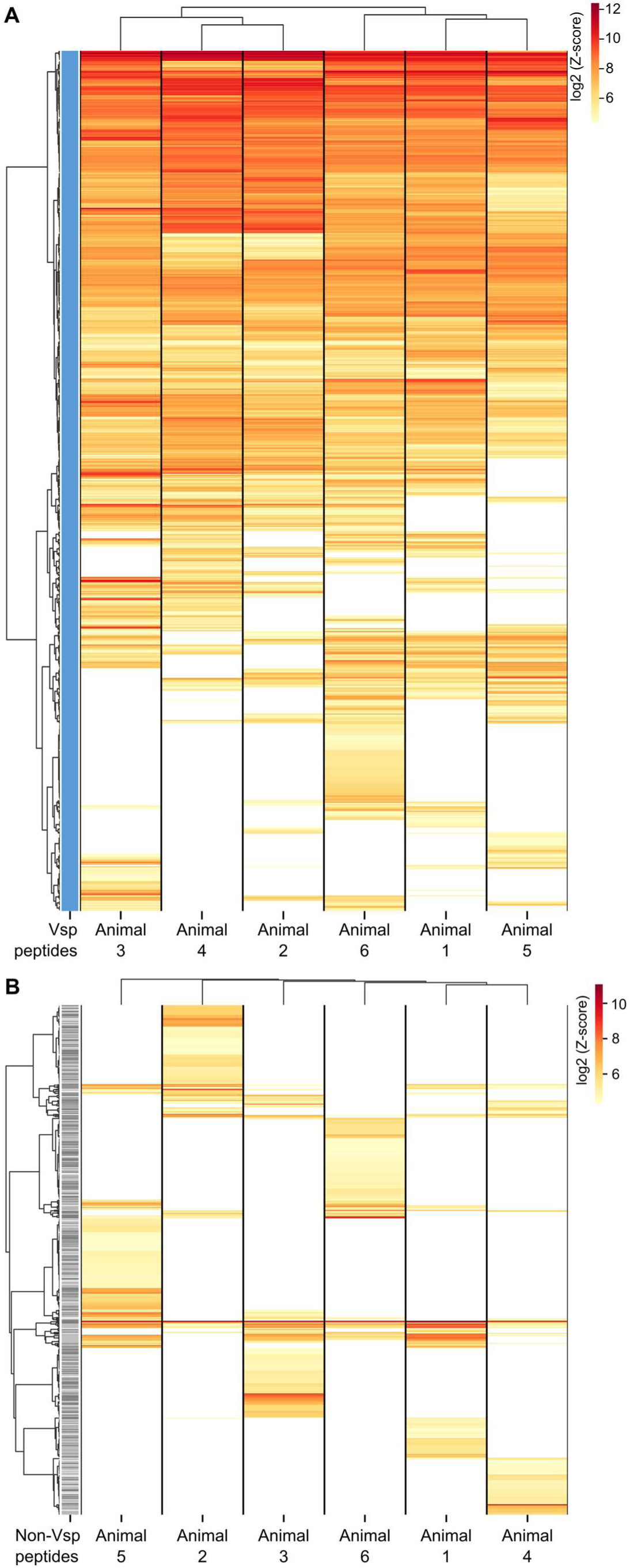
Comparison of peptide enrichment across the six animal samples at week 26. A. Heatmap of the enriched peptides in the 6 animal samples. The Z-score tables were filtered to keep only the peptides enriched in at least one sample and carrying a “Vsp-like” functional annotation. For any given peptide, Z-score values <20 were arbitrarily set to 1, to enable log conversion of the data. Hierarchical cluster analysis was performed on both the samples and peptides, using Ward’s method and Euclidian distance. B. Heatmap of the enriched peptides in the 6 animal samples. The Z-score tables were filtered to keep only the peptides enriched in at least one sample and carrying a “Other” functional annotation. For any given peptide, Z-score values <20 were arbitrarily set to 1, to enable log conversion of the data. Hierarchical cluster analysis was performed on both the samples and peptides, using Ward’s method and Euclidian distance.

**Figure S8:**
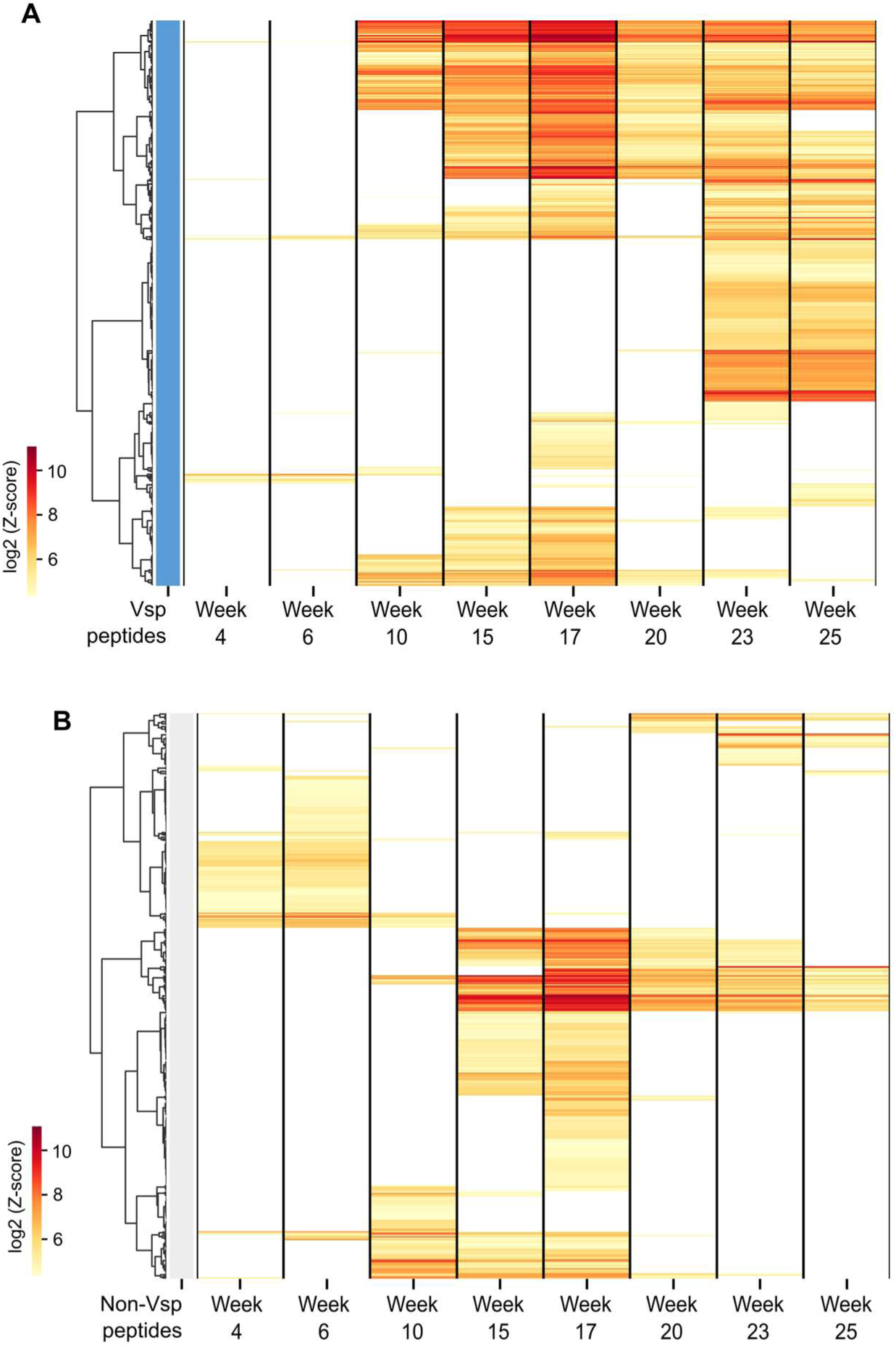
Comparison of peptide enrichment in a single animal during 25 weeks. A. Heatmap of the enriched peptides in the 8 samples of the time-series. The Z-score tables were filtered to keep only the peptides enriched in at least one sample and carrying a “Vsp-like” functional annotation. For any given peptide, Z-score values <20 were arbitrarily set to 1, to enable log conversion of the data. Hierarchical cluster analysis was performed on both the samples and peptides, using Ward’s method and Euclidian distance. B. Heatmap of the enriched peptides in the 8 samples of the time-series. The Z-score tables were filtered to keep only the peptides enriched in at least one sample and carrying a “Other” functional annotation. For any given peptide, Z-score values <20 were arbitrarily set to 1, to enable log conversion of the data. Hierarchical cluster analysis was performed on both the samples and peptides, using Ward’s method and Euclidian distance.

## Notes

https://zenodo.org/records/22749183

### Competing Interest Statement

The authors have declared no competing interest.

